# Fast and accurate resolution of ecDNA sequence using Cycle-Extractor

**DOI:** 10.64898/2026.03.10.710955

**Authors:** Mahsa Faizrahnemoon, Jens Luebeck, King L. Hung, Suhas Rao, Gino Prasad, Ivy Tsz-Lo Wong, Hitesh Kandarpa, Matthew G. Jones, Paul S. Mischel, Howard Y. Chang, Kaiyuan Zhu, Vineet Bafna

**Affiliations:** Department of Computer Science & Engineering, UC San Diego, La Jolla, California 92093, USA; Department of Neuroscience, Scripps Research, La Jolla, California 92037, USA; Department of Biomedical Informatics, Harvard Medical School, Boston, Massachusetts 02115, USA; Department of Pathology, Stanford University School of Medicine, Stanford, California 94305, USA; Indian Institute of Technology, Chennai 600036, India; Department of Biology, Massachusetts Institute of Technology, Cambridge, Massachusetts 02139, USA; Koch Institute for Integrative Cancer Research, Massachusetts Institute of Technology, Cambridge, Massachusetts 02139, USA; Institute for Medical Engineering and Science, Massachusetts Institute of Technology, Cambridge, Massachusetts 02139, USA; Sarafan ChEM-H, Stanford University, Stanford, California 94305, USA; Amgen Research, South San Francisco, California 94080, USA; School of Computer Science, Shanghai Jiao Tong University, Shanghai 200240, China; Halicioğlu Data Science Institute, UC San Diego, La Jolla, California 92093, USA

## Abstract

Extrachromosomal DNA (ecDNA) plays a key role in cancer pathology. EcDNAs mediate high oncogene amplification and expression and are associated with worse patient outcomes. Accurately determining the structure of these circular molecules is essential for understanding their function, yet reconstructing ecDNA cycles from sequencing data remains challenging. We introduce Cycle-Extractor (CE) for this task. CE takes as input a breakpoint graph derived from either short or long read sequencing data and extracts cycles of maximum length-weighted copy number. CE utilizes a mixed-integer linear program (MILP) and a separate traversal procedure, enabling fast optimization and compatibility with open-source ILP solvers.

We evaluate CE against CoRAL (long-read, mixed-integer quadratically constrained programming), Decoil (long-reads), and AmpliconArchitect (AA; short reads) on both simulated data and real cancer cell lines. On simulated ecDNA, CE achieves accuracy comparable to CoRAL across three metrics and consistently outperforms AA and Decoil. On cancer cell lines, CE produces longer and heavier cycles than AA, and performs comparably to CoRAL, while being on average 40× faster. These results demonstrate that CE accurately reconstructs ecDNA from both short and long-read sequencing data, and that long-read inputs allow CE to recover more complete, higher-confidence ecDNA structures. CE improves the prediction of many ecDNA structures. On PC3 ecDNA containing *MYC*, CE uses ONT data to reconstruct a substantially larger and higher-copy sequence (4.2 Mbp) compared to the short-read-derived reconstruction (690 kbp). CRISPR-CATCH experiments confirm the presence of a large ecDNA molecule, validating the long-read-based CE reconstruction.

## Introduction

Extrachromosomal DNA (ecDNA) is the most common source of amplified oncogenes across human cancers (Bailey et al. 2024; Turner et al. 2017; Verhaak et al. 2019; Yan et al. 2024). Studies on large cohorts have revealed that ecDNAs are present in approximately 14%-17% of tumors across multiple cancer types, with particularly high prevalence in glioblastoma and other aggressive malignancies (Bailey et al. 2024; Kim et al. 2020). These circular DNA molecules, typically ranging from kilobases to megabases in size (Bailey et al. 2024), drive tumor formation (Luebeck et al. 2023), evolution (Carroll et al. 1988; Lange et al. 2022; Noorani et al. 2025; Turner et al. 2017) and therapeutic resistance (Haber et al. 1981; Nathanson et al. 2014; Pal Choudhuri et al. 2024). Unlike chromosomal amplifications, ecDNAs lack centromeres and segregate asymmetrically during cell division (Hung et al. 2024; Sankar et al. 2026), leading to rapid changes in copy numbers and intratumoral heterogeneity (Lange et al. 2022). The unique properties of ecDNAs, including their ability to amplify oncogenes (Bailey et al. 2024; Kim et al. 2020), enhance chromatin accessibility (Wu et al. 2019), and facilitate long-range regulatory interactions (Yost et al. 2025), together make them powerful mediators of tumor adaptation and progression.

EcDNAs are not the only circular DNA species in cells. The similarly named extrachromosomal, circular DNA (eccDNA) are smaller circles (1-50 kbp) found at low copy number in all cells (Gao et al. 2024). The vast majority (97%) of eccDNA are smaller than 25 kbp, formed by simple circularization, and do not contain intact and active genes. EccDNA exploration tools focus on the question of presence/absence and quantity (how many molecules can be detected) (Gao et al. 2024; Henriksen et al. 2022; Kumar et al. 2020; Mann et al. 2022; Prada-Luengo et al. 2019; Wanchai et al. 2022; Wang et al. 2021; Zhang et al. 2021). In contrast, we focus here on ecDNAs, which are are much larger, often multi-chromosomal, carry enhancers and intact genes, and are found exclusively in cancer cells (Kim et al. 2020).

To harness the clinical potential of ecDNA, two distinct, albeit related capabilities are needed: *detection* and *reconstruction* of ecDNA sequences. Determining whether or not the tumor genome has ecDNA can provide prognostic information for treatment. Clinical trials are underway for drugs that specifically treat patients with ecDNA (https://clinicaltrials.gov/study/NCT05827614), and detection of ecDNA is required for patient selection. Moreover, large landscape pan-cancer studies correlate ecDNA presence with other phenotypic variables, providing insight into the phenotypic consequence of ecDNA during tumor progression (Bailey et al. 2024; Kim et al. 2020). Reconstruction of the ecDNA sequence, on the other hand, is necessary to understand the content of ecDNA. The ecDNA sequence information has been used to identify oncogenes amplified by ecDNA (Kim et al. 2020), immunomodulatory genes (Bailey et al. 2024), ecDNA with only regulatory sequences (Bailey et al. 2024), ecDNA-ecDNA interaction (Hung et al. 2021), enhancer hijacking (Helmsauer et al. 2020), and the 3-dimensional structure of ecDNA (Chowdhury et al. 2026). We focus here on the reconstruction problem.

Whole genome sequencing (WGS) based approaches have become the standard method for both detection and reconstruction problems. The current computational methods typically follow a parallel strategy for both, starting first with construction of a *breakpoint (or amplicon) graph* that represents amplified genomic regions and their novel adjacencies, using mapped WGS reads (Deshpande et al. 2019; Hadi et al. 2020). For detection, algorithms identify ecDNA by searching for cyclic paths in the graph that exhibit high copy number amplification (Kim et al. 2020; Luebeck et al. 2024).

Because the graph encodes all breakpoints participating in the ecDNA, the reconstruction problem corresponds to extraction of (potentially non-simple) cycles from the amplicon graph. For reconstruction, these cycles must be accurately extracted from the graph to determine the complete ecDNA sequence and structure. Therefore, unlike diploid genome assembly, ecDNA reconstruction faces the following unique challenges. First, ecDNAs can exhibit high structural complexity, containing many breakpoints that create complex rearrangements (Shoshani et al. 2021; Singer et al. 2000). Some breakpoints can be missed by current sequencing and mapping technologies, particularly when they overlap with low-complexity or centromeric regions (Chapman et al. 2023; Keskus et al. 2025; Li et al. 2025; Zhu et al. 2024). Second, large genomic segments can be duplicated within a single ecDNA molecule, appearing multiple times, with potentially different orientations and contexts, which complicates the determination of the correct order of segments (Chowdhury et al. 2026; Hung et al. 2021; Zhu et al. 2024). Third, ecDNA typically occurs as a heterogeneous population - multiple distinct ecDNA species often coexist within the same tumor (Hung et al. 2022). When these species share genomic segments, accurate reconstruction and copy-number estimation become challenging.

Recent developments focus on reconstruction utilizing long-reads (ONT, PacBio) (Giurgiu et al. 2024; Zhu et al. 2024). Long-reads improve cycle extraction by (a) better mapping of all break-points, including those that span repetitive or low-complexity regions, and (b) occasionally spanning regions of high multiplicity reducing ambiguities in reconstruction. CoRAL (Zhu et al. 2024) specifically focused on the problem of extracting cycles that best explain the total copy number and breakpoints in the amplicon graph (see Results), and showed reasonable results on simulations and on cancer cell-lines. However, it uses a mixed integer quadratically constrained programming (MIQCP) that is computationally demanding, and lowers utility for users with no access to MIQCP solvers. Here, we develop Cycle-Extractor(CE), which transforms all quadratic constraints to Mixed Integer Linear Programming (MILP) formulations, improving running time by an order of magnitude while maintaining or exceeding its performance. We tested CE on extensive simulations and on cell-lines, and also tested its ability to detect heterogeneity, where multiple ecDNAs share genomic regions. Finally, by abstracting and separating cycle extraction from graph reconstruction, we could apply CE to better predict ecDNA sequences using only short-read data, increasing its utility for analysis of complex rearrangements.

## Methods

### A high-level description of Cycle-Extractor

Cycle-Extractor (CE) takes an *amplicon graph* (or *CN-weighted breakpoint graph*) and a set of *subwalk constraints* as input (Figure 1), both defined below. The (undirected) amplicon graph is described by *G* = (*V, E, C, ℓ*), where the edge set *E* = *E*_*s*_ ∪ *E*_*d*_ ∪ *E*_*c*_ is the union of *sequence, concordant*, and *discordant* edges. Sequence edges correspond to unbroken chromosomal segments in ecDNA. Discordant edges connect the beginning/end of one sequence edge to itself (indicating *inverted duplication* or *foldback* ), or another sequence edge located elsewhere in the genome; while concordant edges connect the end of one sequence edge to the beginning of an adjacent sequence edge on the same chromosome. Each edge (*u, v*) is associated with a copy-number *C*_*uv*_, and length *ℓ*_*uv*_. The length of the segment edge is the number of nucleotides in the corresponding segment, while *ℓ*(*u, v*) = 0 for all (*u, v*) ∈ *E*_*c*_ ∪ *E*_*d*_.

**Figure 1:**
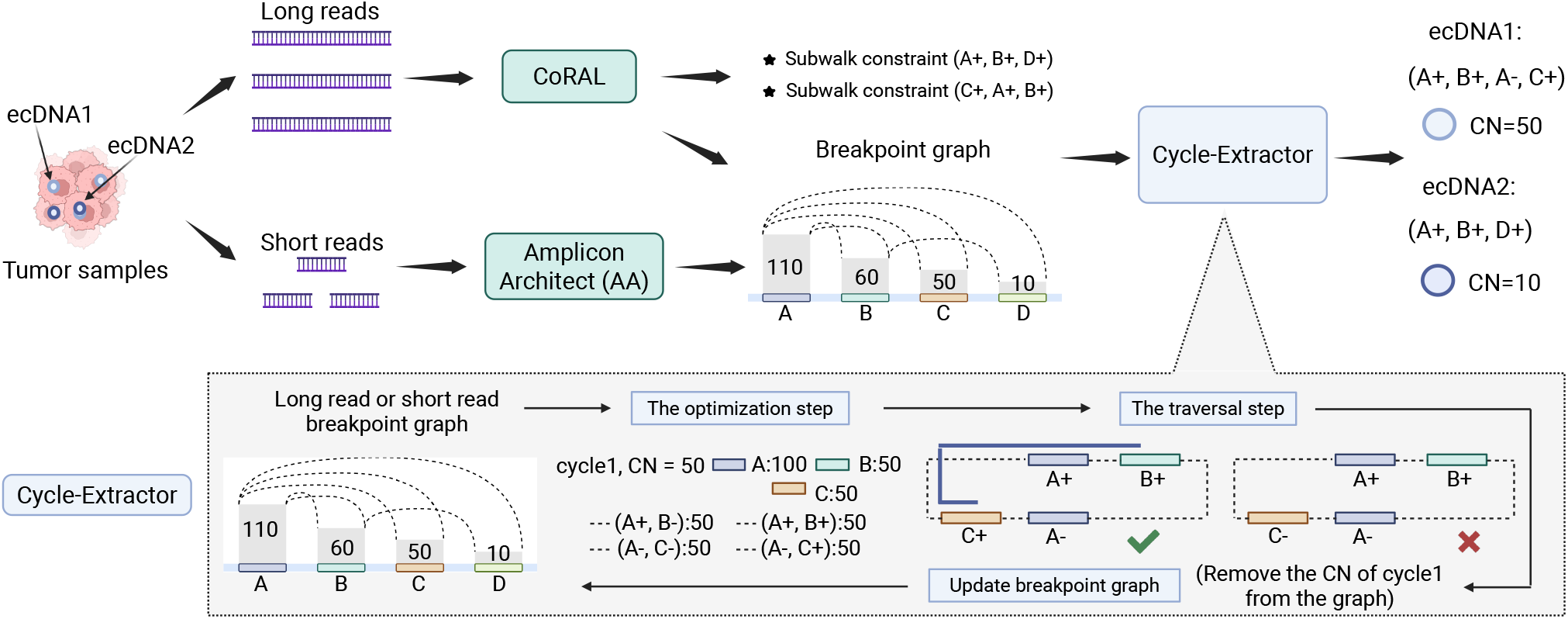
Overview of Cycle-Extractor(CE). CE takes as input, an amplicon graph generated using short-reads (e.g., from AA) or long-reads (e.g., from CoRAL). It applies a mixed-integer linear program to iteratively extract heaviest cycles from the graph (optimization step), and producing ordered traversals of edges (traversal step). From these traversals, CE selects the cycle that satisfies the maximum number of subwalk constraints. The graph is updated by decreasing the copy number of the edges in the selected cycle, and the process is repeated until the amplification is explained.

CE only allows *walks W* in *G* in which segment edges alternate with discordant or concordant edges. A walk *W* is cyclic if it begins and ends at the same vertex. A walk *W* with copy-number *C*_*W*_ is *feasible* if the copy number of each edge (*u, v*) used in the walk satisfies *C*_*W*_ · *m*_*uv*_ ≤ *C*_*uv*_, where *m*_*uv*_ is the number of times an edge (*u, v*) is used in *W* (i.e., its *multiplicity* ). The *length-weighted-copy-number* of a walk is defined by LWCN(*W, C*_*W*_ ) = *C*_*W*_ · length(*W* ). CE optimizes over feasible cyclic walks to output the *heaviest* (maximizing LWCN(*W, C*_*W*_ )) cyclic walk and maximize the number of subwalk constraints satisfied. At a high level, CE has two steps. In the first, *optimization* step, it iteratively computes *C*_*W*_ and an Eulerian subset of edges (*u, v*), along with their multiplicities, that comprise the heaviest cyclic walk^1^. Then the copy numbers from these edges are removed from the graph, and the next iteration proceeds until most of the length weighted copy number the graph *G* (≥ *α* · LWCN(*G*), by default we set *α* = 0.9) is consumed. In the second, *traversal* step, it identifies the actual ordering of the edges in each walk.

One key advantage of long reads is that they provide subwalk constraints that guide reconstruction. Long reads sampled from a cyclic walk *W* are subwalks of *W* . However, real samples may exhibit heterogeneity of structure, where multiple ecDNA may have similar but non-identical sequences, and subwalks may be sampled from different cyclic walks. We model subwalks using 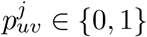 indicating if edge (*u, v*) is in subwalk *P*_*j*_. We then incorporate constraints corresponding to each subwalk. Given a set of subwalk constraints, *P*, CE finds a feasible alternating cycle that simultaneously maximizes LWCN and the number of satisfied subwalk constraints.

### The optimization step of CE

The CE MILP utilizes input encodings and a small number of tunable parameters:

*M* : maximum multiplicity allowed for concordant and discordant edges

*γ*(*W* ): = 0.01 Σ_(*u,v*)∈*W*_ *C*_*uv*_*ℓ*_*uv*_, and balances weights of subwalk constraints relative to LWCN in the objective.

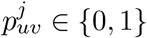: indicates if edge (*u, v*) is in subwalk *p*_*j*_

#### Variables

*F* ∈ ℝ: copy number assigned to edges in the solution but with multiplicity one. *x*_*uv*_ ∈ {0, 1} : *x*_*uv*_ = 1 if edge (*u, v*) is selected, and 0 otherwise.

*f*_*uv*_ ∈ ℝ : copy number assigned to edge (*u, v*) in the solution, should be multiples of *F* .

*y*_*uvm*_ ∈ {0, 1} : (auxiliary) *y*_*uvm*_ = 1 iff (*u, v*) has multiplicity *m* ∈ {1 ≤ *m* ≤ *M* } for each discordant edge in a feasible walk.

*z*_*uvm*_ ∈ ℝ : The auxiliary variable is used to constrain *z*_*uvm*_ = *m* · *F* · *y*_*uvm*_ which is a quadratic formula but is linearized later.

*P*_*j*_ ∈ {0, 1} : indicates if subwalk *p*_*j*_ is satisfied 1 ≤ *j* ≤ *p*.

*d*_*u*_ ∈ {0, 1, 2, ..} : indicates the order of the nodes, used in connectivity constraints.

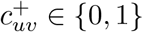: be a binary variable that is 1 if the direction is from *u* to *v*, again used in connectivity constraints.

#### CE MILP

At each iteration, CE maximizes the sum of LWCN and *γ*-weighted subwalk constraint using

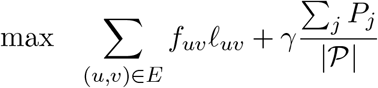

while satisfying the following constraints.

1. **capacity**: For any edge (*u, v*) in a feasible solution, its copy number (*f*_*uv*_) must not exceed its capacity (*C*_*uv*_). If the edge is not chosen in the solution, its copy number is zero.

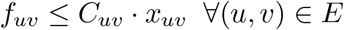
2. **balance**: The sum of the solution copy number of discordant and concordant edges that connect to a node is equal to the solution copy number of the sequence edge connected to that node.

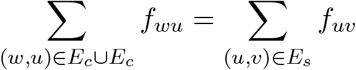
3. **copy number** *F* **with multiplicity one**: Note that if edge (*u, v*) is picked in the solution (i.e., *x*_*uv*_ = 1), then *F* ≤ *f*_*uv*_, but otherwise (*u, v*) imposes no constraint. For this purpose we introduce a sufficiently large constant *C*_max_ (e.g., *C*_max_ = *M* · max_(*u,v*)∈*E*_ *f*_*uv*_).

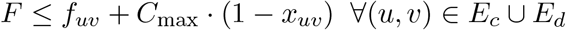
4. **discordant edge multiplicity constraint**: The multiplicity of a discordant edge is managed using

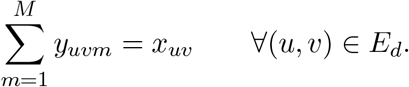

Subsequently the copy of number of the edge is given by the (quadratic) constraint

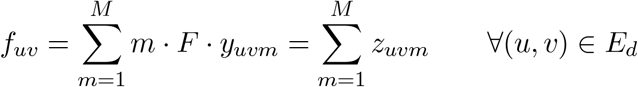

which guarantees *f*_*uv*_ are multiples of *F* . Due to the balance condition, we can prove that *f*_*uv*_ is multiple of *F* for each concordant and each sequence edge (Supplementary Methods).
5. **subwalk constraints**: If a subwalk is in the solution (*P*_*j*_ = 1), all of its edges are in the solution. If a subwalk in not in the solution (*P*_*j*_ ≠ 0), there is not constraint here.

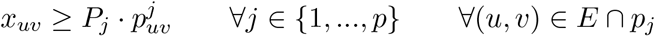

where 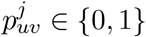 is a constant indicating if edge (*u, v*) is in subwalk *p*_*j*_.

Additional constraints including the linearization of discordant edge multiplicity constraint can also be found in Supplementary Methods.

### Resolving multiple disconnected cycles

CE maximizes the length-weighted-copy-number, aiming to identify the heaviest solution. With no connectivity requirements, it can generate multiple disconnected cycles within a single iteration, as long as LWCN improves. However, this behavior can result in a reduction of the copy numbers for some cycles. For example, if there are two disjoint cycles with length 100 kbp each, and CN 20 and 40, respectively, CE could report a ‘single’ solution outputting both cycles with CN 20 (LWCN=4M). To prevent this, we implemented two methods.

In the first, *connectivity-enforcement* method, we added MILP constraints to enforce that exactly one cycle with maximum LWCN is output per iteration (Supplemental Methods). In the previous example, the constraints ensure that only one cycle with CN=40 is output. The second cycle and other cycles are output one at a time in future iterations. In the second, *CN-enhancement* method, we allowed the MILP to output multiple disjoint cycles initially, but subsequently increase the CN of each cycle to the minimum copy number of any edge in that cycle. Among all cycles initially produced, we output the cycle with the maximum enhanced copy number, and reiterate. In the Results section below, we refer to connectivity-enforcement method as CEc, and the CN-enhancement method as CE, treating it as the default. We tested CE and CEc for accuracy and speed.

### The traversal step of CE

The output of the optimization step is a set of edges, (*u, v*) ∈ *W*, and the multiplicities assigned to them, 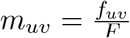 . The graph defined by these edges is Eulerian and may admit multiple Eulerian cycles (Brightwell and Winkler 2004). We perform multiple (default 100) traversals and select a cyclic walk which satisfies the maximum number of subwalk constraints. A modified Hierholzer’s algorithm is used to perform the traversal and is applied as follows:

CE maintains a count *r*_*uv*_ of the number of times each edge (*u, v*) remains unused, initialized as its multiplicity *m*_*uv*_. It starts with the sequence edge (*u, v*) ∈ *E*_*s*_ with positive *r*_*uv*_ that begins at the canonically smallest chromosome and position, with positive orientation (i.e., starting with the node corresponding to the smaller position). The algorithm then alternates between selecting a non-sequence edge (discordant or concordant) and a sequence edge. When a non-sequence edge is to be chosen, CE randomly selects a discordant or concordant edge (adjacent to the current node) with *r*_*uv*_ *>* 0 and heads to the next node. Since each node connects to only one sequence edge, the next sequence edge is automatically determined; and if the node corresponds to the larger endpoint in the sequence edge, the orientation is negative, otherwise it is positive. This process continues until the walk returns to its starting node, thereby forming a closed walk. Finally, all closed walks are iteratively merged into a single walk by identifying shared nodes and rotating the edges for the cycle to be inserted.

## Results

### Datasets and metrics

We simulated 75 distinct cyclic structures using ecSimulator (https://github.com/AmpliconSuite/ecSimulator) closely following CoRAL (Zhu et al. 2024). Very briefly, each structure contains a collection of between 1 and 20 oriented genomic segments joined end to end. The end of the last segment is joined to the beginning of the first segment to complete the cyclic structure. The simulated structures modeled the known mechanisms of ecDNA formation including episomal (Carroll et al. 1988), chromothripsis (Ly et al. 2019; Shoshani et al. 2021), and 2-foldback (Brewer et al. 2011; Passananti et al. 1987), and were additionally modified using internal structural variations. Next, long and short reads were simulated from the simulated ecDNA sequence at one of three coverages (50X, 100X, or 250X) modeling the increased copy number of ecDNA, and merged with reads from one of five simulated normal, diploid genomes (each with 13X coverage).

From long-read input, an amplicon graph was generated using CoRAL. Cycle reconstruction was performed using CoRAL, Cycle-Extractor and Decoil (Giurgiu et al. 2024) for comparisons. For short-reads, the amplicon graph was generated using AA, and cycle extraction was performed using AA and Cycle-Extractor.

We additionally analyzed 31 amplicon graphs from cancer cell-lines (Supplementary Table S1, 26 with Illumina WGS, and 12 amplicons with long-reads (ONT) whole genome sequencing, with both sequencing modalities for 7 amplicons. All cell-lines are known to carry ecDNA from prior cytogenetics experiments. For the cell-lines, the true ecDNA sequence is not known, and therefore, we tested them on the previously defined LWCN (length-weighted-copy-number) and LWCNR (ratio) metrics. The tools are all designed for the user to have some control on the stopping criteria. For all experiments below, and all methods (except AA), we stopped when 90% of the total LWCN in the graph was explained, or if the current iteration had not solved the MILP after 2 hours. For AA, such a control is not available, and the default mode was used.

### CE matches or exceeds CoRAL, Decoil and AmpliconArchitect (AA) performance in simulations

No single metric captures the quality of reconstructed cycles, and we utilized 3 metrics: First, to measure the nucleotide overlap, consider the true and reconstructed cycles each as multi-sets *T, R* (including multiplicities) of nucleotide positions. The **Cycle Interval Overlap (CIO)** metric given by

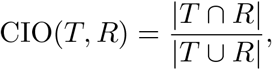

represents the Jaccard index of the shared nucleotides. Next, **Reconstruction Length Error (RLE)** reports the normalized difference in amplicon lengths computed as

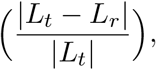

where *L*_*t*_ and *L*_*r*_ denote the lengths of the true and reconstructed cycles, respectively. Finally, **Cyclic Longest Common Subsequence (LCS)** measures the length of the longest common subsequence contained in the true cycle and the reconstructed cycle after eliminating intervals that are not found in both, and normalizing to the length of the true cycle.

CE performed at least as well as CoRAL (Figure 2A–C) and outperformed Decoil across all three metrics. For the CIO metric, Decoil was correct about 40% of the time in contrast with CE and CoRAL, which were correct 80% of the time. In about 20% of the samples, Decoil did not return any cycles. CE improved upon CoRAL for the LCS metric in nearly half of the cases. A partial explanation could be better satisfaction of subwalk constraints, where CE satisfied 7.6% more subwalk constraints than CoRAL. Additionally, CoRAL makes a design choice which gives the same priority to extracting walks and cycles. In some cases where a breakpoint edge is missed, CoRAL selects a walk that is heavier than any available cycle. In summary, the CoRAL and CE reconstructions, while not identical, match up well overall.

**Figure 2:**
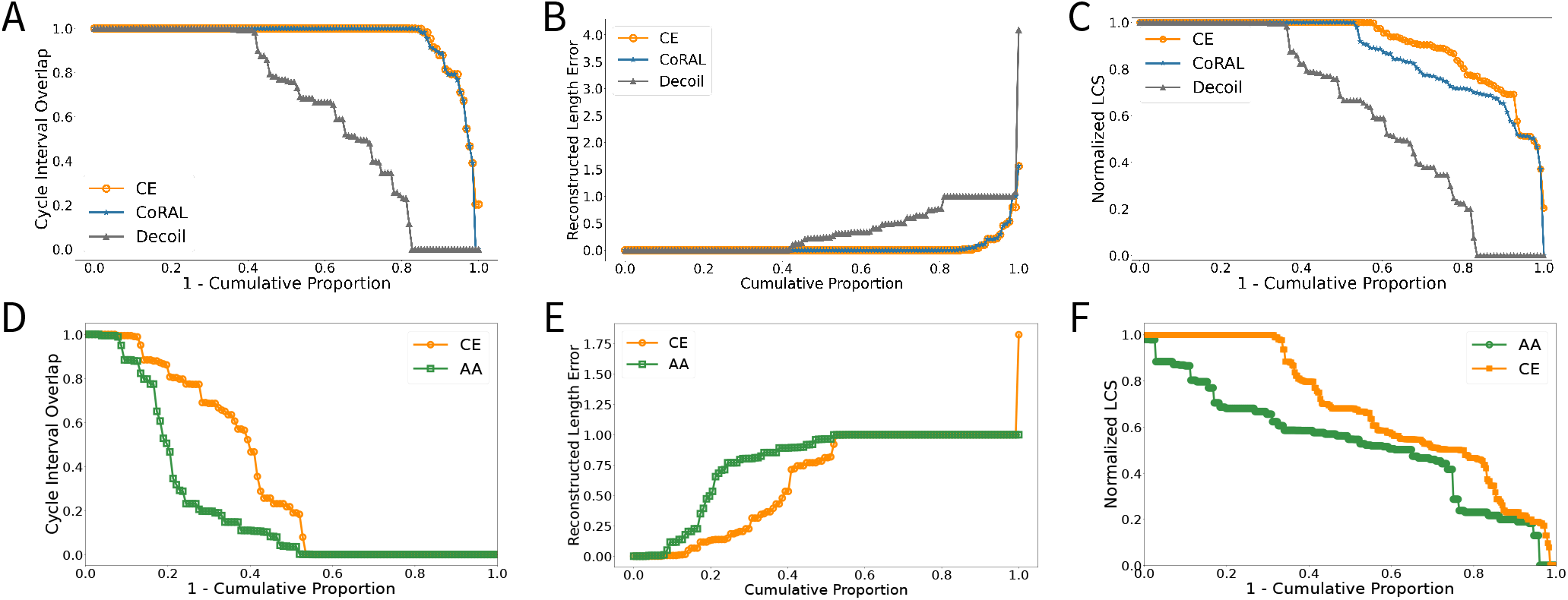
CE accuracy in reconstruction on simulated data. (A-C): CIO, RLE, and LCS metrics comparing CE to CoRAL and Decoil on long-read simulations. For any threshold of a metric the results show the fraction that matched or improved upon the metric. (D-F): Comparison between CE and AA on short-read simulations.

The cyclic LCS metric could be dominated by a single long, unbroken segment even when the breakpoint ordering elsewhere was incorrect. To assess if this inflated our LCS metrics, we plotted the distribution of segment lengths and ecDNA sizes (Supplementary Figure S1) . A large majority of the fragments were less than 250 kbp, much smaller than the average size of ecDNA (2Mbp), discounting long fragments as impacting the score. Moreover, CE had 100% accuracy in 62% of the cases, which can be achieved only with a perfect reconstruction.

To assess the stability of the traversal step, we performed multiple runs of the randomized Eulerian traversal on the optimized edge sets. We evaluated the consistency of the resulting cycle orderings using the normalized longest common subsequence (LCS) between cycles obtained across runs. The inferred cycles exhibit high agreement across runs (Supplementary Figure S2) . Importantly, we did not observe substantially different ecDNA structures across runs. While minor variations in traversal order may arise due to local ambiguities in the Eulerian graph, these differences do not affect the overall structure or the set of constituent edges in the inferred cycles. This stability is expected, as the MILP step significantly constrains the solution space, leaving limited flexibility in the traversal.

For short-reads, CE performed much better than AA on all metrics, showcasing the power of optimization in extracting the heaviest cycle (Figure 2D-F) . While the amplicons used to simulate short and long-read data were the same, the long-read data missed fewer breakpoints and also provided subwalk constraints. Therefore, the cumulative performance of CE on short-reads was worse than its own performance on long-reads across all three metrics (Supplementary Figure S3). Manual exploration confirmed that this was due to missed breakpoints in short-read analysis, showcasing the additional sensitivity of long-reads for breakpoint detection. In some simulations, missed breakpoint edges prevented cycle formation (CIO=0 and RLE=1) or shortened the cycle. This happened often with short-reads but sometimes also for long-reads.

Finally, we also tested CE on additional difficult cases where two true cycles sharing genomic segments were simulated. When the two cycles had very similar copy numbers, CE merged them into a longer, heavier cycle (Supplementary Figure S4 A). However, when the copy numbers were very different, CE predicted the two cycles accurately (Supplementary Figure S4 B). Heterogeneity of ecDNA is not well studied and CE provides a method for its exploration in real data.

#### CE reconstructs heavier cycles in cancer cell-lines

To test performance on cell-lines where ecDNA have been previously reported, we utilized whole genome ONT reads and CoRAL to generate an amplicon graph for 12 ecDNA amplicons. We compared the cycle extractions of CE, CE with connectivity-enforcement (CEc), and CoRAL on the amplicon graphs. Because the cycle reconstructions may differ and produce different orderings, we chose the sum of the length weighted copy number ratio (LWCNR) for the three heaviest cycles identified by each method. The sum LWCNR describes how much of the copy number of the amplified region is explained by the three heaviest cycles.

CE, CEc and CoRAL exhibited nearly identical behavior (Figure 3A), both selecting cycles with high LWCNR as their top-ranked cycles, and generally explaining a large fraction of the amplicon. SNU16 was the only exception where CEc and CoRAL each returned a single cycle, whereas CE extracted 5 cycles covering a larger share of the LWCN. The two stopped for different reasons. CoRAL reconstructed one cycle and reported the remainder as paths; CEc found no additional cycle or path within the per-iteration time limit and terminated after the second iteration. Notably, the one (heaviest) cycle extracted by CEc was identical to the heaviest cycle extracted by CE. In some other samples, CoRAL obtained lower LWCNR because it prioritized a heavy walk rather than a cyclic structure as the dominant amplicon. When considering all metrics, the total copy number, length, and LWCNR, comparing the three heaviest cycles is not meaningful, and we tested those for the single heaviest cycle. Here also, CE, CEc and CoRAL cycles matched up well (Figure 3B). In contrast, when taking a short read amplicon graph as input, AA generally produced cycles with lower LWCNR (mean 0.598) compared to CE (mean 0.704) (Figure 3C,D). The performance of CE and CEc was effectively identical across all samples.

**Figure 3:**
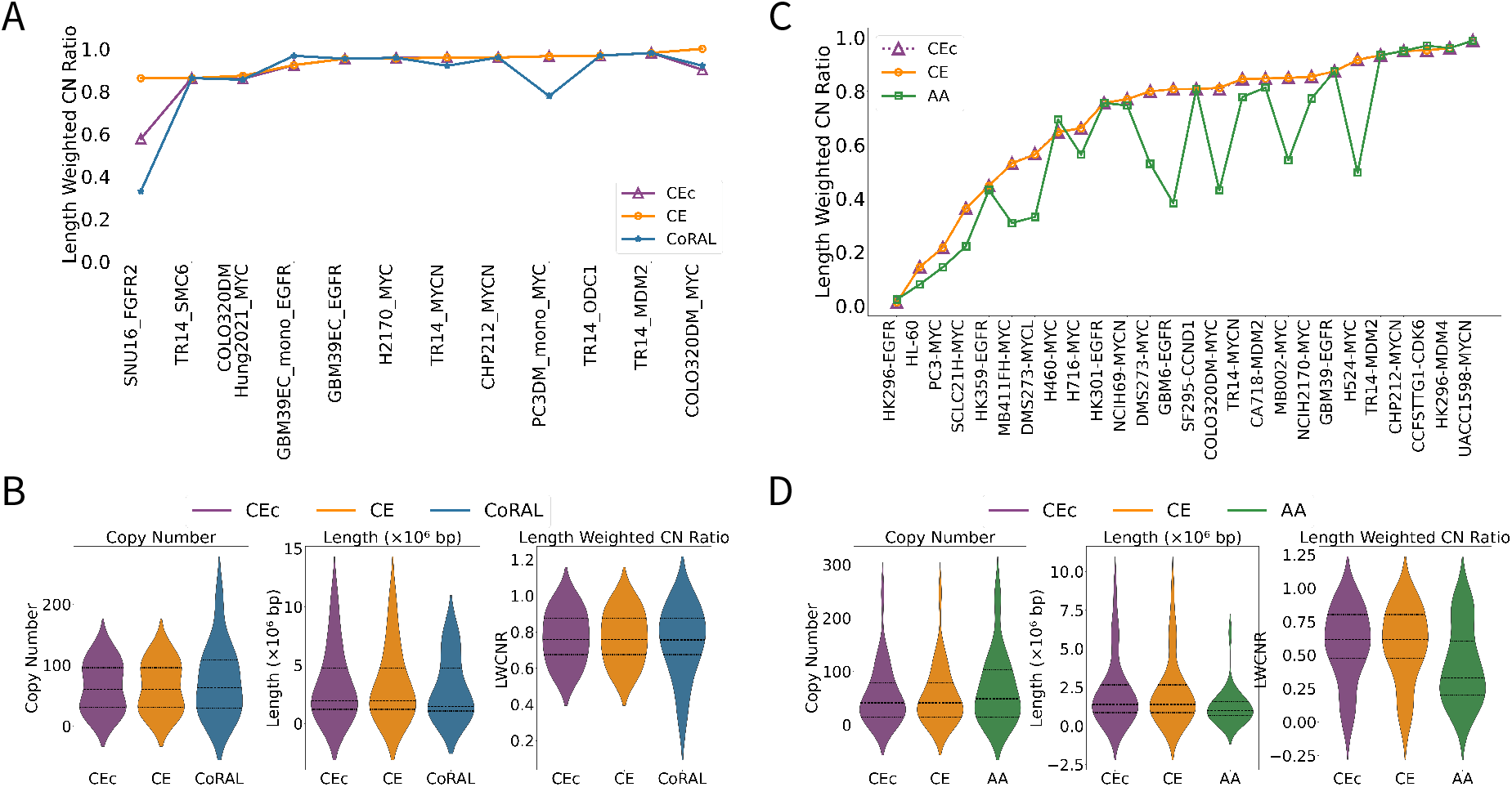
CE performance on cancer cell-lines. (A) Comparison of LWCNR of the three heaviest cycles extracted by CE, CEc and CoRAL on ONT sequenced cell-lines. (B) Copy Number, length and LWCNR of the heaviest cycle on ONT derived graphs. (C) LWCNR of the 3 heaviest cycles extracted by CE, CEc and AA on Illumina sequenced cell-lines. (D) Copy Number, length and LWCNR of the single heaviest cycle on Illumina derived graphs.

We also compared the performance of all methods using cell-lines for which both ONT and Illumina sequencing was available. Here the copy number estimations in these biological replicates are comparable but not identical across platforms; and the short read and long read amplicon graphs may differ in segmentation and extent, so an aggregate LWCNR comparison would not reveal how the reconstructions themselves differ. We therefore inspected the LWCN, length, and CN of the cycles produced for each cell line individually (Supplementary Figures S5-S9). In cases where the graph had a relatively simple structure (e.g., the CHP-212 ecDNA amplifying *MYCN* ), the cycles extracted by CoRAL and CE using the ONT graph and by AA and CE using the Illumina graph were all identical. In other cases, the cycles extracted by CoRAL and CE using the ONT amplicon graph were heavier and longer than the cycles extracted by AA and CE using Illumina. Moreover, even on the Illumina amplicon graph, the CE extracted cycle was heavier and longer than the AA extracted cycle.

Decoil could not be included in this comparison (Figure 3A,B), because it does not report copy numbers for its reconstructed cycles. We therefore compared it on length alone, against the heaviest cycle extracted by each of the other methods. Decoil recovered an identical cycle in a few cell lines, but in others its cycles were substantially shorter, and for several amplicon graphs none of the Decoil cycles overlapped the graph’s segments at all (Supplementary Figure S10).

As an additional test, we also evaluated tools that were designed for detecting smaller circular eccDNA, which are often *<* 10 kbp, and ubiquitously found in normal and cancer cells (Gao et al. 2024). Specifically, we tested CReSIL (Wanchai et al. 2022), which showed superior performance compared to other eccDNA tools (Gao et al. 2024). In a small test on 2 simulated samples (where ground truth ecDNA structure was known), CReSIL did not find any ecDNA. On the cell-line CHP212, where a single ecDNA is known (Chamorro González et al. 2023; Helmsauer et al. 2020), CReSIL missed the ecDNA but identified 115 smaller eccDNA non-overlapping with the single ecDNA.

Together, these results indicate the power of using long-reads and a principled cycle extraction algorithm for exDNA reconstruction.

### CE runs 40× faster than CoRAL

Because CE is a cycle extraction method, and uses the graph reconstruction of CoRAL, we first compared cycle extraction running times. On both simulated data and cell-lines, CE was faster than the other methods. For the simulated benchmark, the running times of CE, CoRAL, and Decoil were all measured using a single thread to enable a fair comparison, as Decoil does not support multi-threading whereas both CE and CoRAL do. On simulated data, the average running time after graph construction was 0.31 seconds for CE, 3.69 seconds for CoRAL, and 53.6 seconds for Decoil.

Decoil was not included in the cell-line comparison, because it reports running time for the entire sample rather than per amplicon, so its runtime is not directly comparable to the per-amplicon extraction times of CE and CoRAL. CE and CoRAL were therefore evaluated using their default multi-threaded implementations. CE completed its runs in 0.13 to 75.47 seconds, with 58% of the samples finishing within one second (Figure 4A). CoRAL required between 0.26 seconds to 32.5 minutes, and its mean running-time was 40× higher than that of CE. CEc ranged from 0.18 seconds to 10 minutes, finishing within one second on 42% of samples. Notably, all cases, with one exception SNU16, terminated without reaching the time limit of 2 hours per iteration. SNU16 was one challenging example with 71 discordant edges and 338 edges in total, where as described above CoRAL completed its execution normally but extracted only one cycle, after which it extracted paths rather than additional cycles. CEc also extracted one cycle in its first iteration but did not identify another cycle or path within the per-iteration time limit in the second iteration. Therefore, the runtime reported for CEc and CoRAL corresponds to the extraction of the one single cycle. CE by contrast, took 8.37 seconds to extract 5 cycles which explained 90.73% LWCN of the graph.

**Figure 4:**
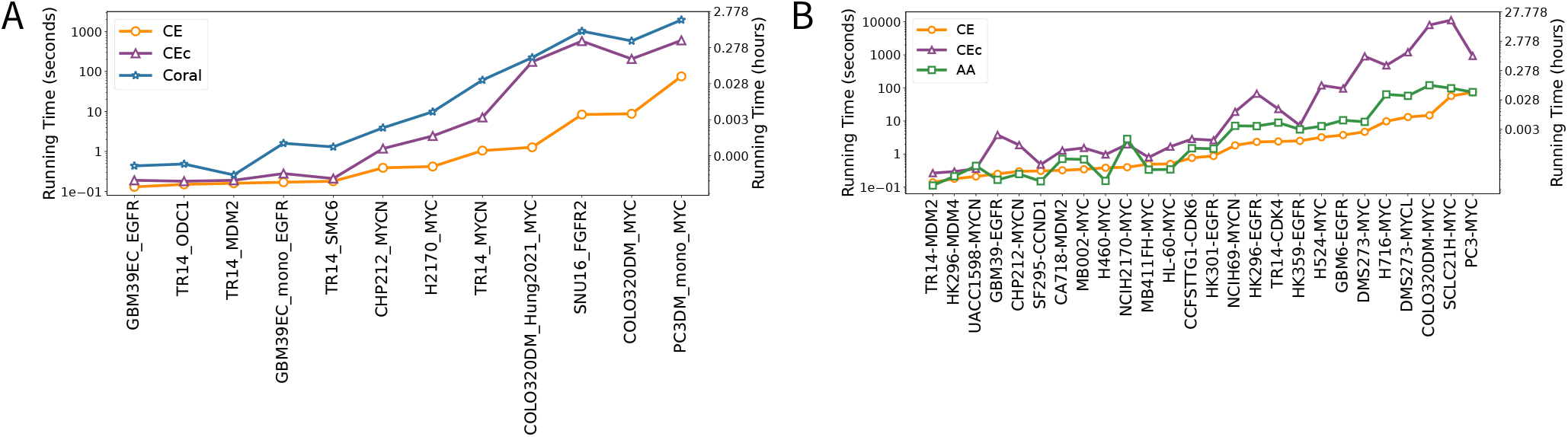
Running times of different methods for cycle extraction. (A) Cycle extraction on ONT samples. (B) Cycle extraction on Illumina samples. All cases, with the exception of SNU16 on ONT reads, terminated without reaching the time limit of 2 hours per iteration. The CoRAL and CEc running time for SNU16 are for extracting a single heaviest cycle.

The results on Illumina short-reads were similar. CE continued to be the fastest method, running faster than AA on 75% of the samples despite the MILP based cycle extraction procedure (Figure 4B), while maintaining much higher reconstruction quality (heaviness). For CEc, application of connectivity enforcement method introduces additional (though, increasing linearly in the graph size) integer variables, increasing CEc running time compared to CE. The running time of CEc was also higher when using the short-read graph compared to its time on the long-read graph. Because of missing breakpoints, the cycles extracted were shorter, and more cycles were extracted, leading to higher running time for CEc.

On both simulated long-read data and cell lines, the running time of CE increased with graph complexity (i.e., the number of edges, Supplementary Figure S11A,B), though it was not the sole determinant. For example, CE resolved the PC3 MYC amplicon with 310 edges (133 sequence, 118 concordant, 59 discordant) in 75 seconds, while the larger SNU16 graph (338 edges in total, 138 sequence, 129 concordant, 71 discordant) was solved in under 10 seconds. This is because CE identified multiple disconnected cycles early in the SNU16 iterations, reaching the termination criterion (90% graph LWCN coverage) more quickly.

To evaluate the practical performance of the complete workflow, we compared the total runtime of graph construction and cycle extraction for CE, CEc, and CoRAL on the long-read cancer cell lines (Supplementary Figure S12). Since CE and CEc use the breakpoint graph generated by CoRAL, graph construction was included in the runtime of all three methods. Over the complete pipeline, CE remained substantially faster than CoRAL, achieving an average 11-fold reduction in runtime. The total runtime of CE ranged from 2.02 to 93.03 seconds, compared with 2.05 to 643.53 seconds for CEc and 3.14 to 1967.56 seconds for CoRAL. Furthermore, 83.3% of the datasets completed in less than one minute using CE, compared with 66.7% for CEc and 58.3% for CoRAL. These results demonstrate that the computational advantage of CE extends beyond the cycle extraction step and translates into a substantial reduction in the runtime of the complete long-read reconstruction workflow.

#### CE reconstructions improve our understanding of ecDNA biology

The algorithmic approach utilized by CE improves upon our ability to reconstruct ecDNA sequences, both on short reads, and also on long-reads where additional subwalk constraints can be satisfied. The tested cell lines included GBM39 where the CE results matched the previously predicted structures exactly (Hung et al. 2021; Wu et al. 2019). However, notable differences were observed between CE and Decoil for three amplicons: the *MYC* amplicon in H2170, and the *ODC1* and *SMC6* amplicons in TR14 (hereafter H2170-*MYC*, TR14-*ODC1*, and TR14-*SMC6* ). For H2170-*MYC*, the Decoil reconstruction was substantially shorter than the corresponding CE and CoRAL cycles, and for TR14-*ODC1* and TR14-*SMC6*, Decoil did not reconstruct any cycle overlapping with the amplicon graph (Supplementary Figure S10). To further validate the cycles produced by CE, we utilized evidence from chromatin-capture (Hi-C) data acquired previously (Chowdhury et al. 2026; Hung et al. 2021). For H2170-*MYC* (Figure 5A), the Decoil reconstruction contained only Chromosome 8 segments, whereas CE (and CoRAL) reconstructed an ecDNA containing segments from Chromosomes 8 and 17. The Hi-C map showed strong interactions between the Chromosome 8 and Chromosome 17 regions, supporting the CE reconstruction and indicating that the Decoil reconstruction was incomplete. For TR14-*SMC6* (Figure 5B), the Hi-C data showed strong interactions between segments from Chromosomes 1 and 2, consistent with the CE reconstruction. A similar result was observed for TR14-*ODC1* (Figure 5C).

**Figure 5:**
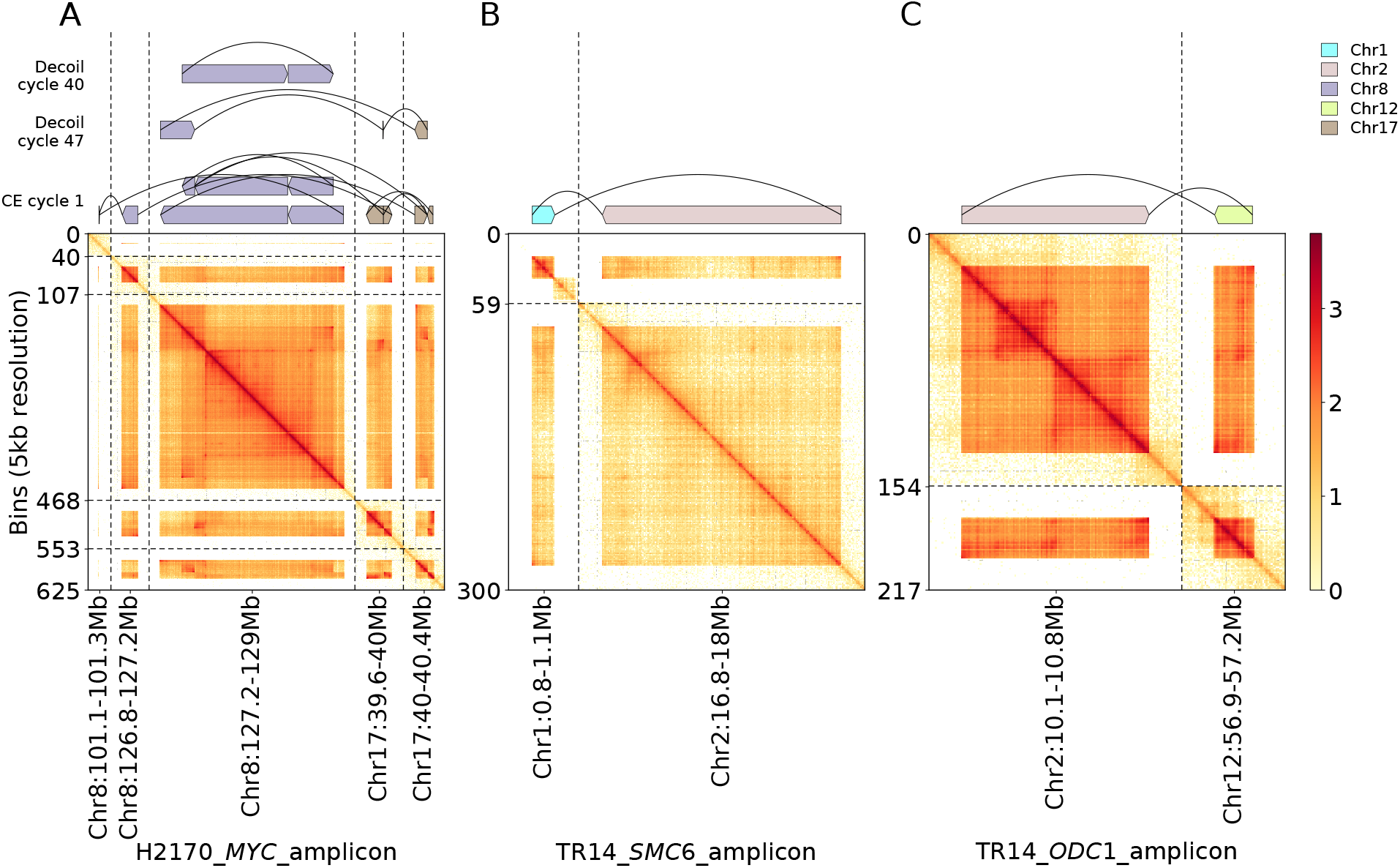
Hi-C evidence for CE reconstructions. (A) H2170-*MYC*, (B) TR14-*SMC6*, (C) TR14-*ODC1*. In each panel the heatmap shows the log_10_ Hi-C contact matrix over the amplified intervals at 5 kb resolution. The track above each heatmap shows the reconstructed structures over the same coordinates, including the heaviest CE cycle (bottom row) and the Decoil cycles overlapping that amplicon graph (rows above).

For Illumina reads acquired from the CA718 cell line, AA predicted three cycles: one carrying *MYCN* and *MDM2*, another carrying *PDGFRA*, and a third non-genic cycle (Figure 6A). In contrast, CE identified two cycles, effectively merging the non-genic cycle predicted by AA with the *MDM2* +*MYCN* cycle (Figure 6B) to generate a longer and heavier cycle. The CE reconstruction is a more parsimonious explanation of the amplicon and suggests that predictions of non-genic ecDNA containing only regulatory sequences (Bailey et al. 2024) should be orthogonally validated. We also observed that the two cycles predicted by CE (and by AA) share genomic segments. Our reconstructions do not preclude the presence of alternative species generated by recombination across the shared fragments.

**Figure 6:**
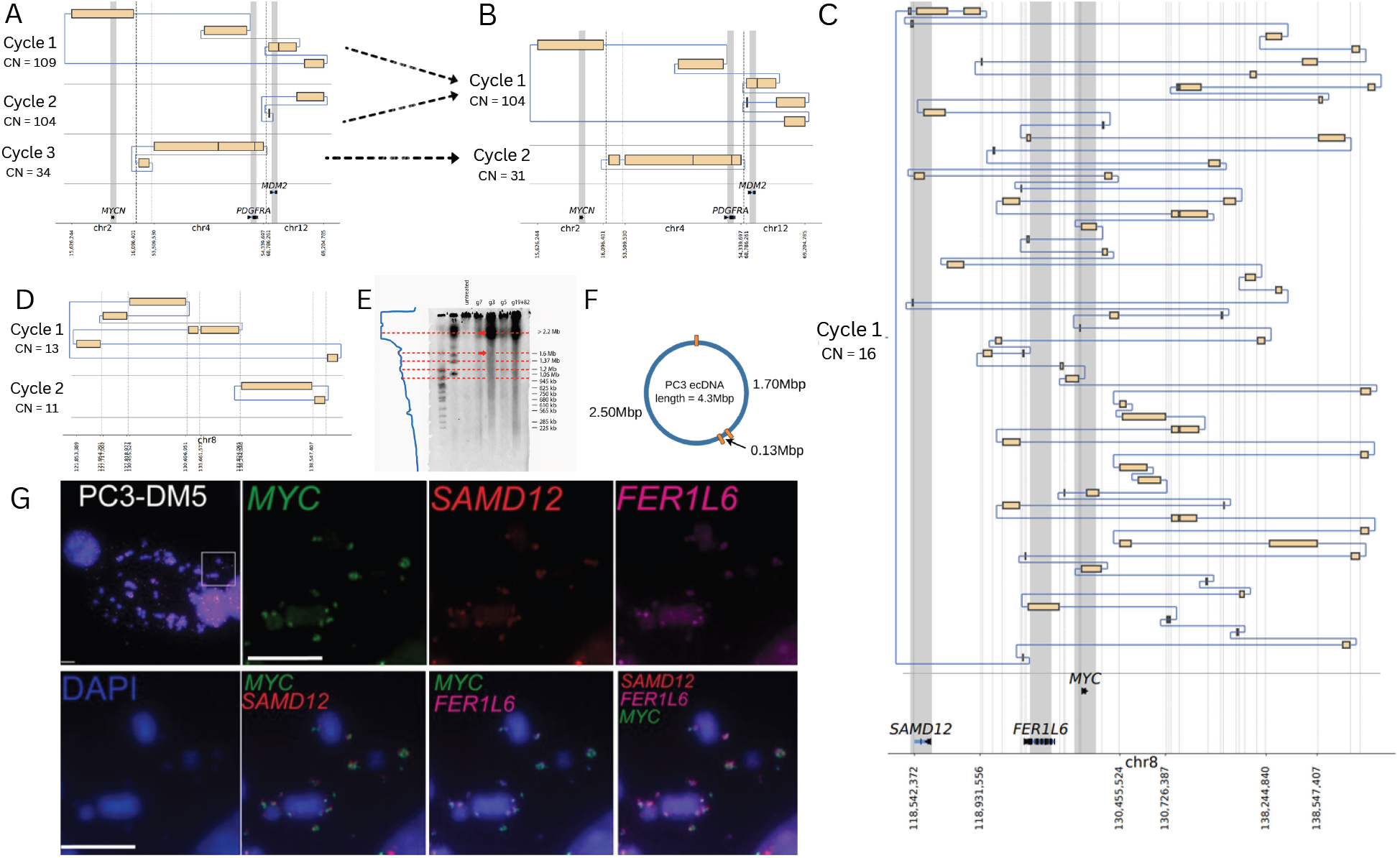
AA and CE reconstructions for specific cell lines. (A) AA reconstruction with three cycles for CA718. (B) CE reconstruction of CA718 ecDNAs, producing two cycles. (C) CE reconstruction of the PC3 amplicon. (D) CE reconstruction of PC3 using the AA (Illumina) graph. (E) CRISPR Catch data for the CE amplicon suggest fragments longer than 2.2 Mbp and 1.6 Mbp. (F) Schematic reconstruction of the PC3 ecDNA with gRNA sites marked. (G) DNA FISH validation using probes for the *MYC, SAMD12*, and *FER1L6* loci. Co-occurrence of the three probes supports the CE long-read reconstruction.

We also compared the CE cycles generated on a PC3 amplicon using Nanopore and Illumina sequencing. The CE cycle reconstructed from Nanopore was substantially larger and more amplified than those obtained from the short-read data, predicting a 4.32 Mbp cycle (Figure 6C) explaining 78% of the amplicon copy number. In contrast, the cycle extracted using Illumina reads (Figure 6D) had length 690 kbp and explained 9% of the amplicon. The CN of the oncogene *MY C* was 12 copies in the short-read cycle versus 47 copies in the long-read cycle. To validate our data, we performed a CRISPR-Catch experiment (Hung et al. 2022) where single CRISPR mediated cuts linearized the ecDNA, and pulse-field gel electrophoresis was used to separate the linearized product. Multiple guide RNAs were used to linearize the structure, and produced fragments with different lengths (Figure 6E). Notably, gRNA g3 produced fragments of lengths *>* 2.2, and 1.7 Mbp. On the CE reconstruction of PC3 ecDNA, g3 occurred thrice, and supported fragments of size 2.5, 1.7, and 0.13 Mbp (Figure 6F), consistent with the CRISPR-Catch data. The gRNA g5 is contained in the oncogene *MYC*. It appeared twice on the cycle and supported fragments of size 0.12 and 4.3 Mbp. The other gRNA were controls which did not appear on the predicted cycle. All of these were consistent with the CRISPR-Catch data.

The CE-reconstructed ecDNA using long-reads included many loci that were not in the shortread reconstruction, for example in regions encoding *SAMD1* and *FER1L6*, while also containing the *MYC* locus that was originally reported for short-read reconstruction (Figure 6C; shaded regions). To validate, we performed a DNA multi-FISH experiment (Supplementary Methods), with probes for the 3 loci. The results showed co-occurrence of the probes in multiple ecDNA (Figure 6C), supporting the reconstruction.

To further characterize the PC3 reconstruction, we analyzed the chromosomal, repeat, satellite, and gene composition of the short-read and long-read PC3(DM) cycles using KaryoScope v2.0.0 (Ranallo-Benavidez et al. 2026). The two reconstructions had highly similar sequence composition (Supplementary Tables S2-S6, comprised largely of Chromosome 8, and similar proportions of repetitive elements (45-48% LINE, with smaller proportions of SINE, LTR, and DNA transposons), though the long-read cycle contained significantly higher genomic content in the longer sequence. The long-read cycle had slightly higher genic content (44% versus 54%; Supplementary Figure S13). Supplementary Figure S13 also shows that the long-read reconstruction extends into genomic regions surrounding *SAMD12* and *FER1L6*, while retaining the *MYC* locus identified in the short-read reconstruction.

### CE is robust to the choice of optimizer

We evaluated CE on its option to run with the HiGHS optimization (Huangfu and Hall 2018) module against the default option to run Gurobi. HiGHS reconstructed the same cycles as Gurobi on all cell lines and on all but one simulated dataset. The single discrepancy was caused by numerical precision, where the HiGHS solution exceeded the graph copy-number constraints by approximately 10^−8^, resulting in a slightly longer but technically infeasible cycle. We also compared the runtime of the two solvers across all cell lines. CE with Gurobi (mean runtime: 7.41 Illumina, 8.04 ONT) was generally faster than with HiGHS (mean:17.07 Illumina, 645.03 ONT), with the largest differences observed on the long-read datasets (Supplementary Figure S14A,B). CE with HiGHS was, however, faster than CoRAL on all evaluated cell lines except PC3DM mono.

## Discussion

Reconstruction of the ecDNA sequence is important to understand the content of ecDNAs identified in cancer genomes. Nevertheless, it remains a very challenging problem for two reasons. First, ecDNA can carry multiple larger regions with higher multiplicity, which is akin to the problem of repeats in genome assembly, but harder because the regions of high multiplicity are very large often exceed 100 kbp.

We describe an algorithmic approach wherein a cycle is extracted from the amplicon graph that best explains the copy number and breakpoints represents the current structure. Unlike previous methods, the CE methods utilizes an MILP optimization that is an order of magnitude faster while maintaining accuracy. Moreover, CE is designed to work off both Illumina and ONT reads and can be easily adapted to other sequencing technology. On long-reads, it additionally optimizes the use of subwalk-constraints for better disambiguation of sequences. For both short and long read, the input amplicon graphs required as input to CE are now readily produced from WGS data using robust available tools (Luebeck et al. 2024; Zhu et al. 2024), making it seamless to use.

In extensive tests, CE reconstructed ecDNA structures with high accuracy. Some metrics such as LWCNR could favor CE, which optimizes length weighted copy number (LWCN). However, when the ground truth (the true cycle) was known, for example in simulated data, we only used metrics that were independent of CE’s objective function, thus providing a fair comparison for all methods.

When ground truth was not known, for example in cell-lines, LWCNR was a reasonable evaluation metric. It did not require access to ground truth data, and higher values suggest that a larger fraction of the amplified region was explained by the predicted ecDNA, providing a parsimonious explanation. For many well studied cell lines, we obtained the identical answer to the previous (often manual) reconstructions from earlier studies. In 3 other scenarios wehre the CE reconstructions were substantially different from Decoil or where Decoil did not return a cycle, chromatin interaction maps supported the ecDNA call showing interactions between different segments that were joined in the reconstruction, but not in their neighborhoods. In one case (PC3DM), where we obtained a substantially novel reconstruction, we performed CRISPR Catch and DNA FISH experiments for orthogonal validation. The results strongly supported the CE reconstructions, and provided support for the CE objective function.

In ongoing future work, we will create modules for AmpliconSuite and CoRAL so that cycle extraction can be automatically performed using CE. By determining the ecDNA sequence, CE will have direct utility for investigating functional aspects of ecDNA including enhancer hijacking (Helmsauer et al. 2020), regulatory rewiring, presence and absence of genes (Bailey et al. 2024; Kim et al. 2020) and other functional elements, the DNA repair mechanisms used (Rose et al. 2025), evolution of ecDNA structure (Luebeck et al. 2023) and others.

The second challenge with ecDNA reconstruction is the heterogeneity where multiple ecDNA share genomic segments, and it is not clear if they represent the same ecDNA or different ones. Notably, those ecDNA could exist and function similarly as a single molecule or multiple molecules, or even both in one sample. In preliminary results, CE can extract two ecDNA sequences with shared segments if they have distinct copy numbers. When distinct ecDNA species share both extensive genomic sequence and have very similar copy numbers, CE merges them into a single cycle. Thus, a subclonal event that generates a new cycle from an existing one will be identified only when the new cycle achieves a very different copy number. Note however, that in the absence of other information, this merged reconstruction represents the most parsimonious explanation of the data. As sequencing technology improves to sequencing longer reads, the additional subwalk constraints will help resolve the correct structure(s). In future work, we will also extend our methods by directly measuring the extent of heterogeneity in terms of conflicting subwalk constraints and the extent of genomic region sharing among multiple cycles that together explain the amplification.

In general, unambiguous determination of the correct ecDNA sequence will be best achieved by a combination of sequencing and other experimental methods. These include single molecule sequencing extensions such as CRISPR-Catch (Hung et al. 2022), single-cell methods like ATAC-seq (Sundaram et al. 2024) and Hi-C (Chowdhury et al. 2026; Zhao et al. 2024) and orthogonal technologies such as DNA-FISH (Luebeck et al. 2024) and others. These methods can be applied to individual samples of interest, and future work will seek to combine these methods with the algorithmic analysis of amplicon graphs. In the meantime, CE can serve as an easy to use and relatively accurate method for ecDNA reconstruction.

## Code Availability

The Cycle Extractor source code is publicly available under an open-source license at: https://github.com/AmpliconSuite/CycleExtractor. The repository includes detailed documentation, example datasets, and scripts for reproducing key results presented in this study. The version of CE used in the presented analysis is also provided as Supplemental Code.

## Acknowledgments

This work was carried out as part of the eDyNAmiC team, supported by the Cancer Grand Challenges partnership funded by Cancer Research UK (CGCATF-2021/100025 [V.B.]) and the National Cancer Institute (OT2CA278635 [V.B.]; U24CA264379, R01GM114362 [V.B.]). K.L.H. was supported by a an NCI Predoctoral to Postdoctoral Fellow Transition Award (NIH K00CA274692) and is a Howard Hughes Medical Institute Fellow of the Damon Runyon Cancer Research Foundation, DRG-2570-26. This material is based upon work supported by the National Science Foundation Graduate Research Fellowship Program under Grant No. DGE-2545911. Any opinions, findings, and conclusions or recommendations expressed in this material are those of the author(s) and do not necessarily reflect the views of the National Science Foundation.

M.F., K.Z., and V.B. conceived the study. M.F. developed the CE formulation, implemented the software, and performed the comparison between CE and other tools. J.L., S.R., and H.K. contributed to software development. M.G.J. generated the simulated data and long-read sequencing of cell lines. J.L. compiled the short-read cell lines for the AmpliconArchitect comparison. K.L.H. performed the CRISPR-CATCH experiments and G.P. analyzed the data. I.T.L.W. performed the DNA-FISH of PC3 and interpreted the results. K.Z., P.S.M., H.Y.C., and V.B. supervised the study. M.F., K.Z., and V.B. drafted the manuscript. All authors read and approved the final manuscript.

## Supplementary Figure Captions

**Supplemental Figure S1:**
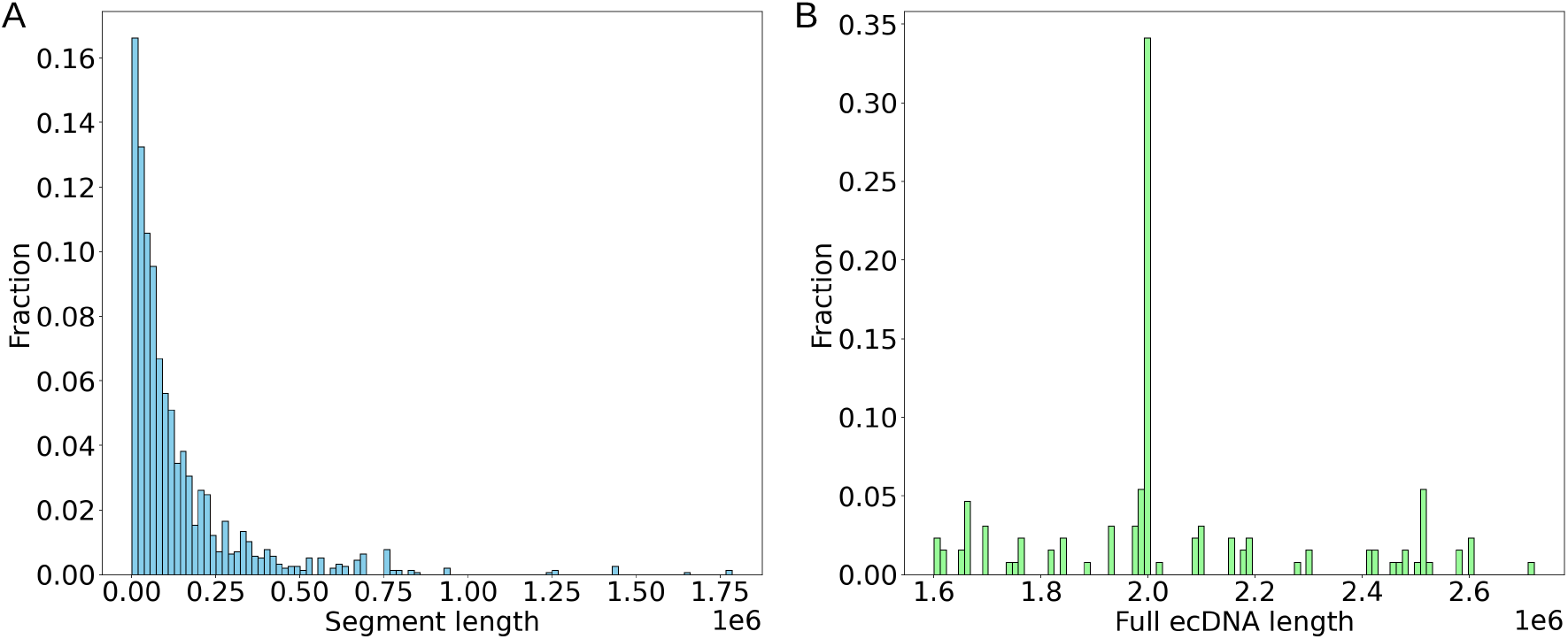
The lengths of the segments versus the lengths of the ecDNA. The large majority of the segments are smaller than 250K, while ecDNA are larger, with average size exceeding 2Mbp.

**Supplemental Figure S2:**
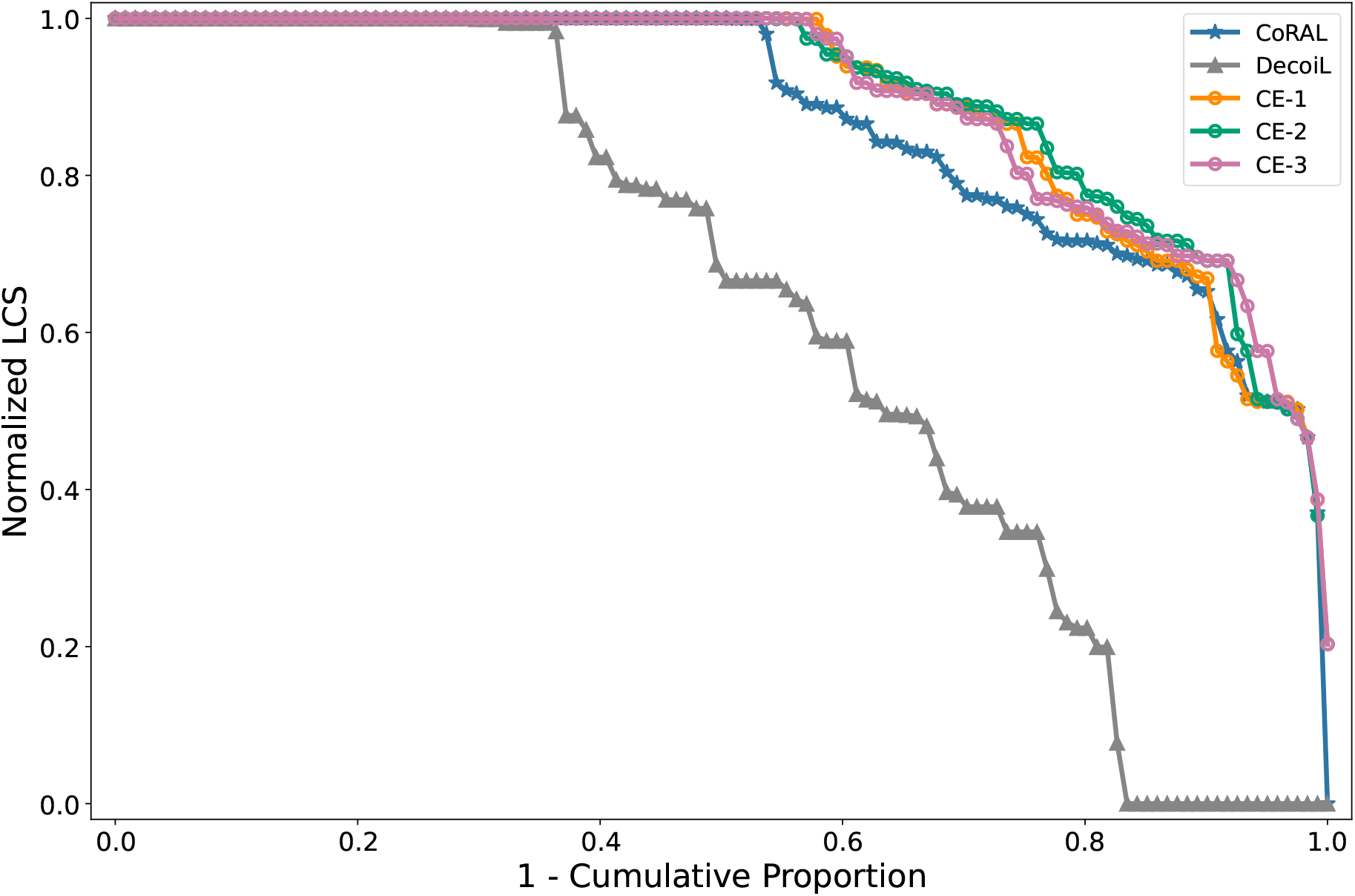
Stability of cycle ordering under randomized traversal. Cumulative distribution of normalized longest common subsequence (LCS) values obtained from three runs of the traversal step with different random orders on the same optimized edge set. Curves CE-1, CE-2, and CE-3 correspond to three independent runs. The LCS metric for CoRAL and Decoil are added for comparison here.

**Supplemental Figure S3:**
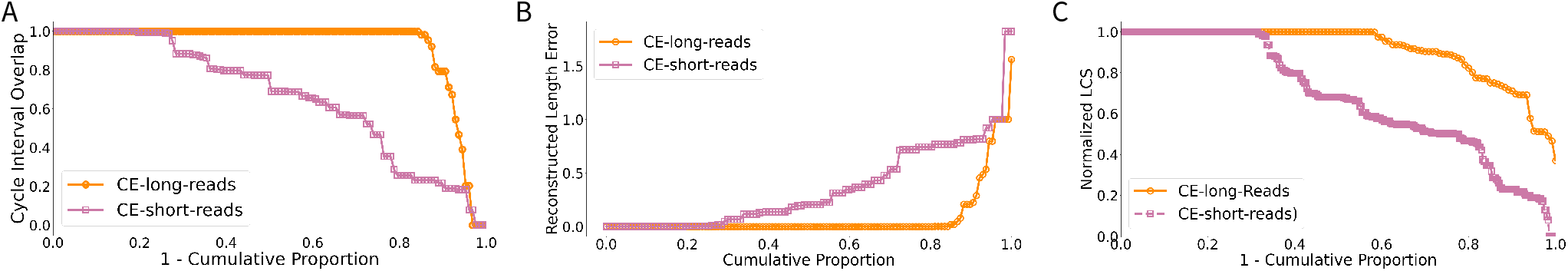
Comparison of (A) CIO, (B) RLE and (C) LCS metrics by CE on simulated data using long-read and short-read graphs.

**Supplemental Figure S4:**
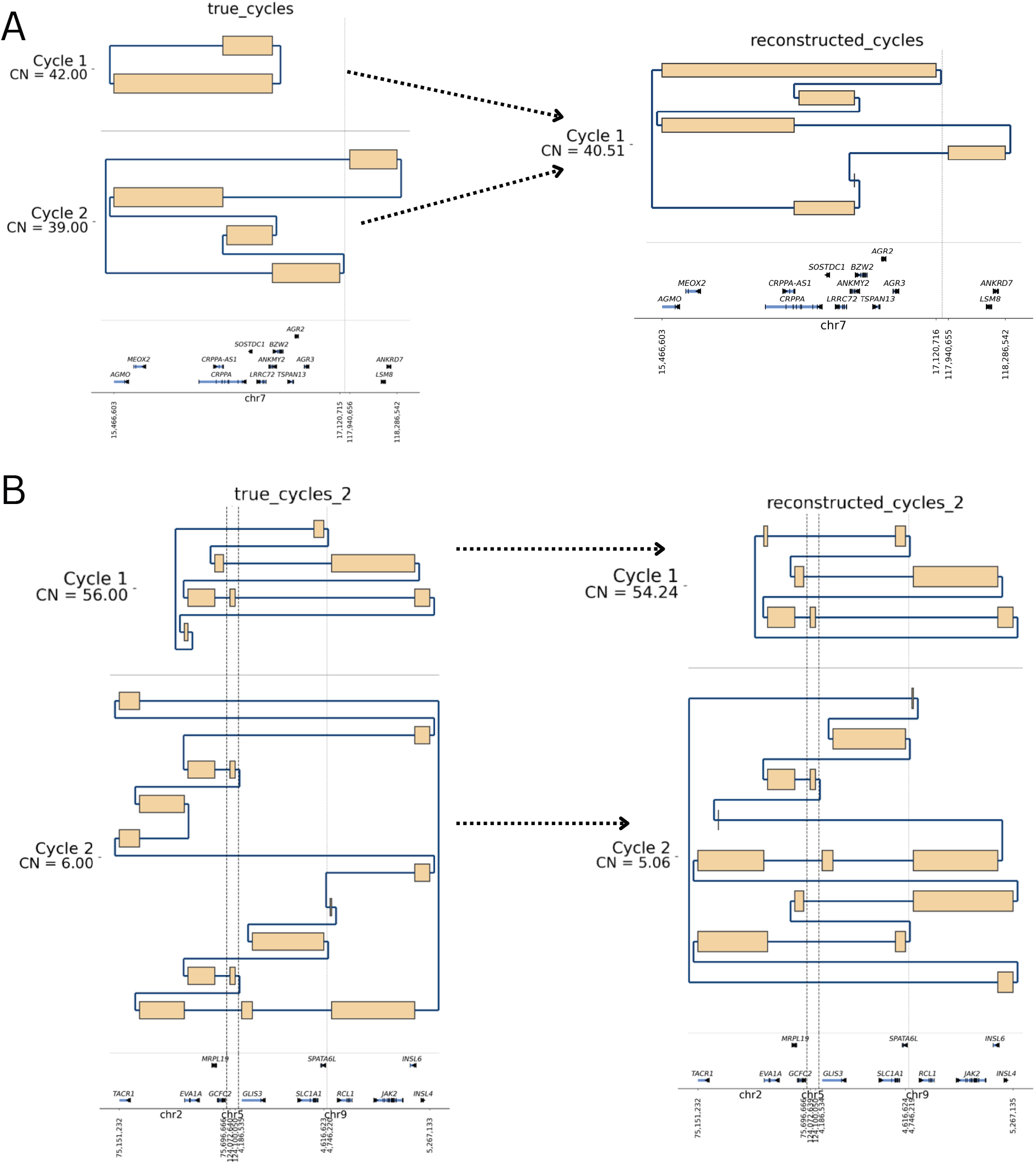
CE behavior on heterogeneous data. (A) CE merges two true cycles onto one when they share genomic segments and are very similar in copy number. (B) CE predicts two separate cycles when the true cycles have very different copy numbers.

**Supplemental Figure S5:**
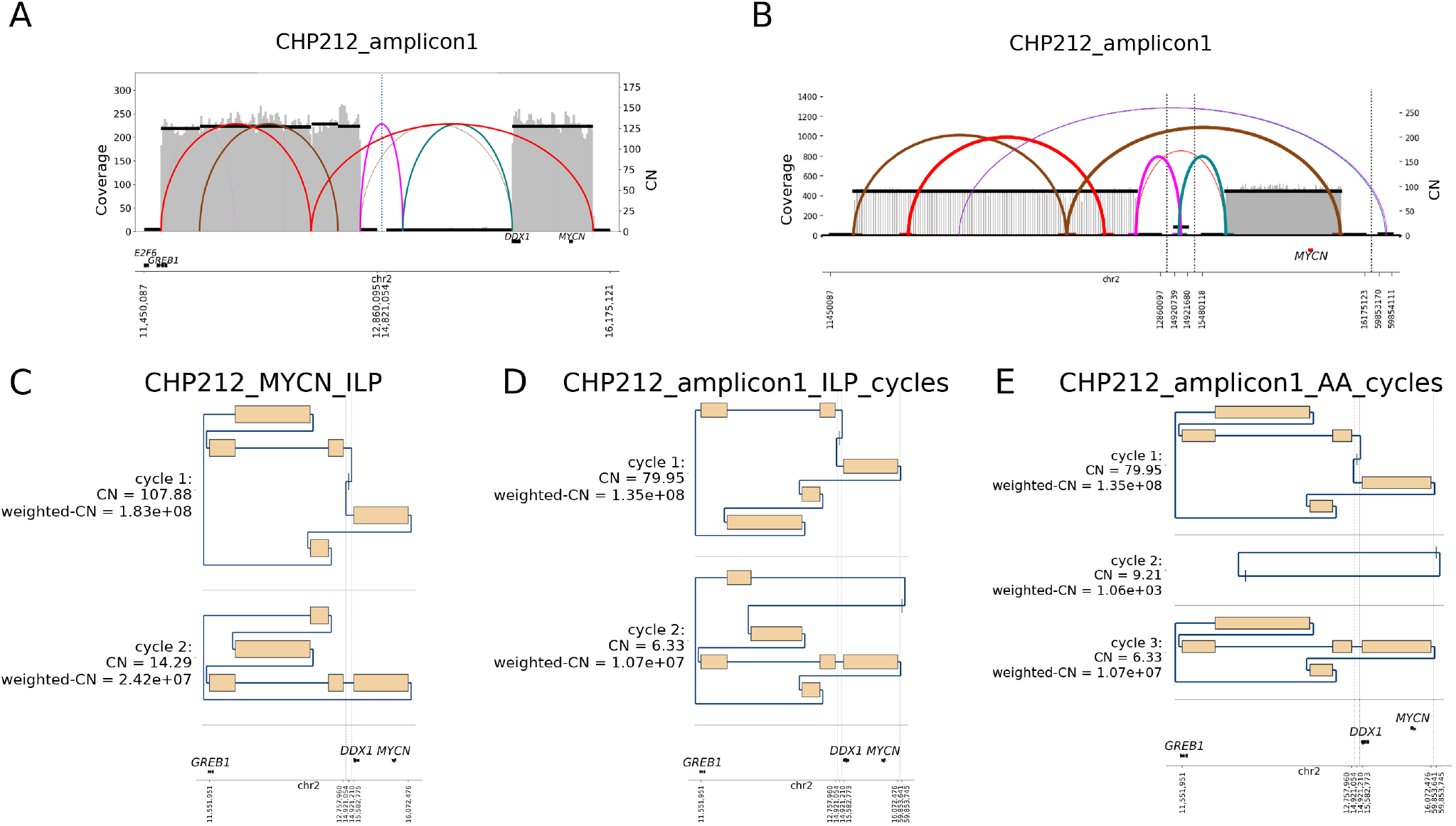
CE reconstruction on CHP-212 cell line. (A) Long read amplicon graph. (B) Short read amplicon graph. (C) CE reconstruction using long-read graph with 2 cycles, which is identical to CoRAL. (D) CE reconstruction using short-read graph. (E) AA reconstruction (AA can only extract cycles from short-read graph) with 3 cycles.

**Supplemental Figure S6:**
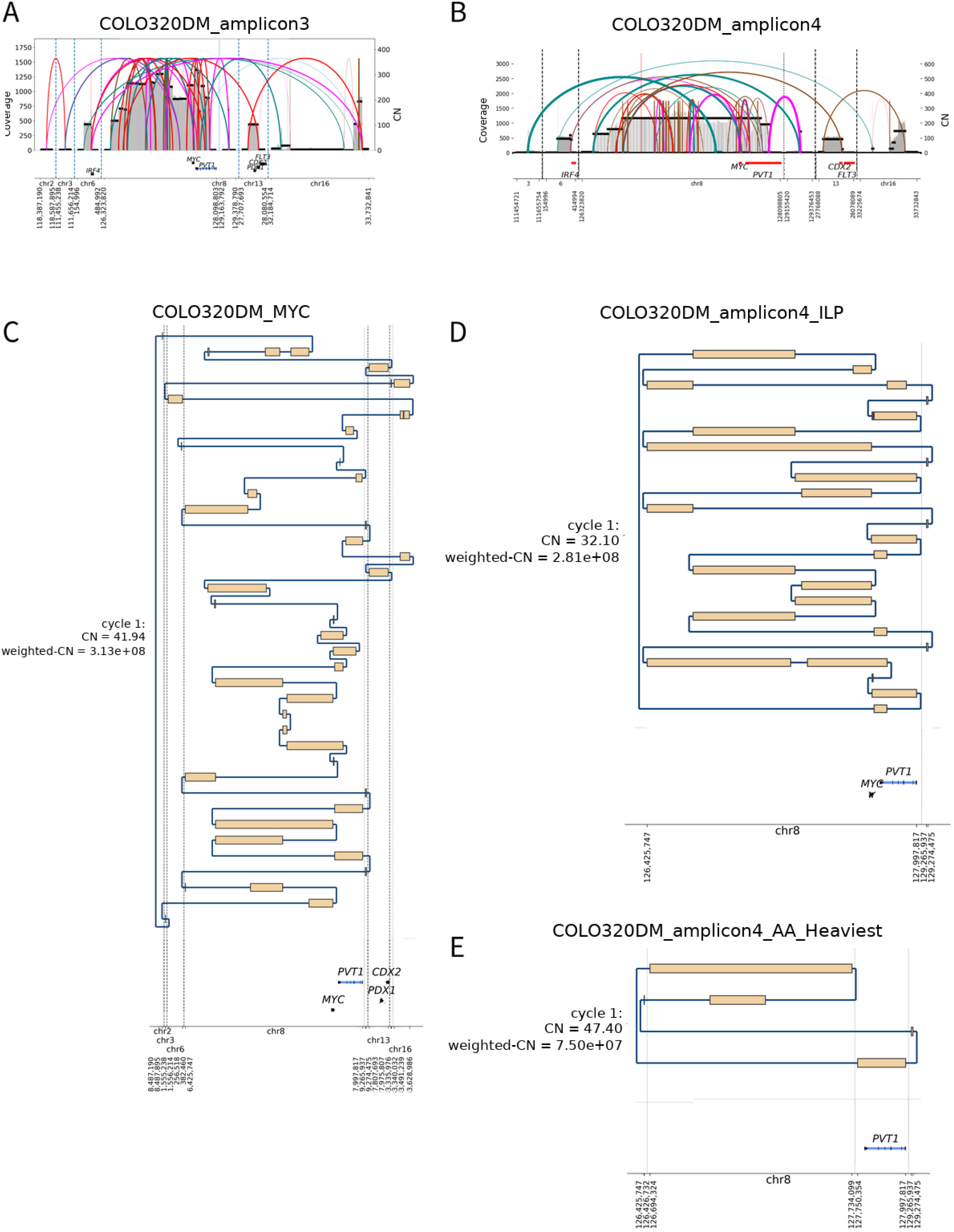
CE reconstruction on COLO320DM cell line. (A) Long read amplicon graph. (B) Short read amplicon graph. (C) The heaviest cycle from CE reconstruction using the long-read graph, which is identical to CoRAL. (D) The heaviest cycle from CE reconstruction using short-read graph. (E) The heaviest cycle from AA reconstruction.

**Supplemental Figure S7:**
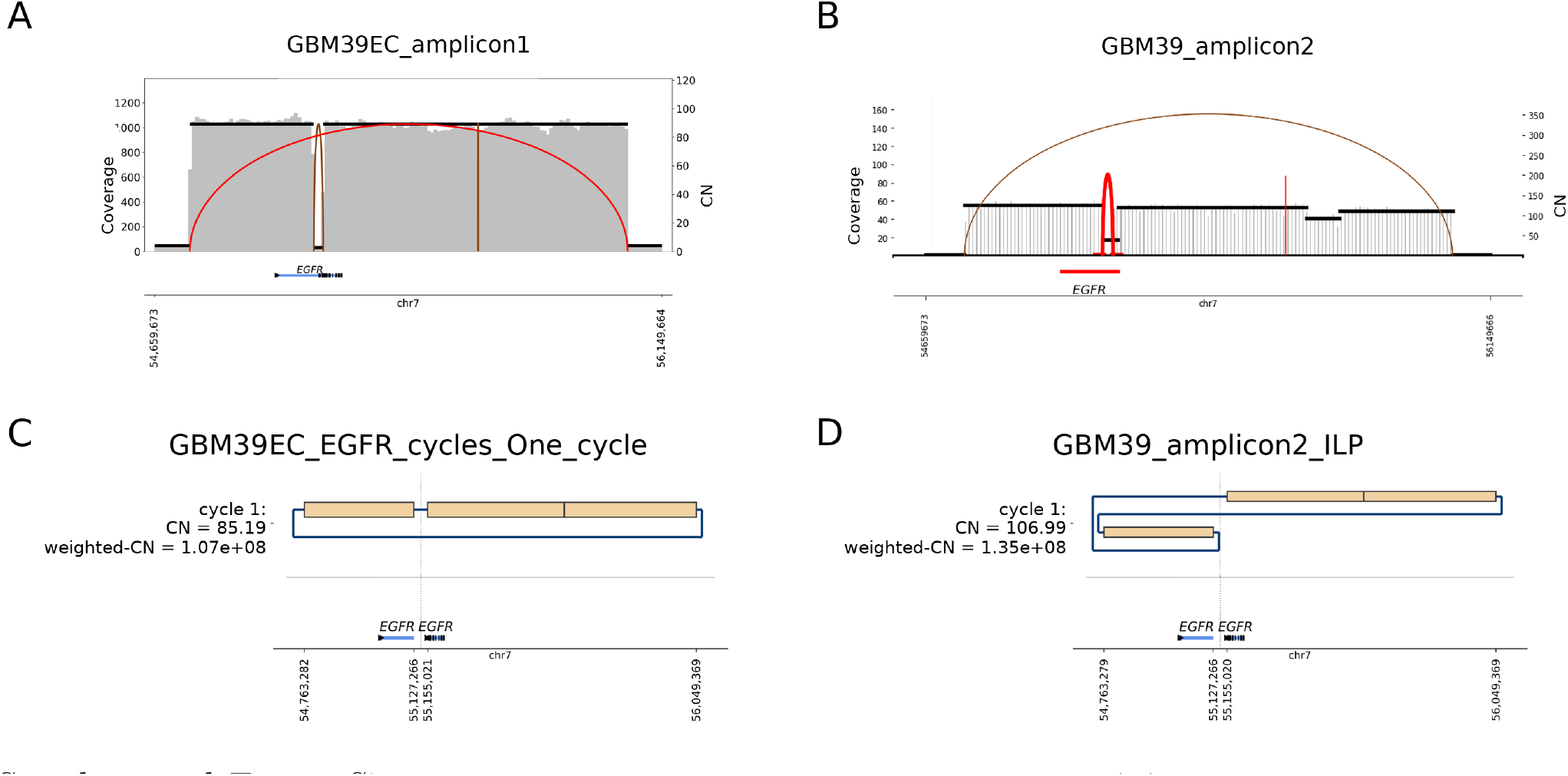
CE reconstruction on GBM39 cell line. (A) Long read amplicon graph. (B) Short read amplicon graph. (C) The heaviest cycle from CE reconstruction using the long-read graph, which is identical to CoRAL. (D) The heaviest cycle from CE reconstruction using the long-read graph, which is identical to AA.

**Supplemental Figure S8:**
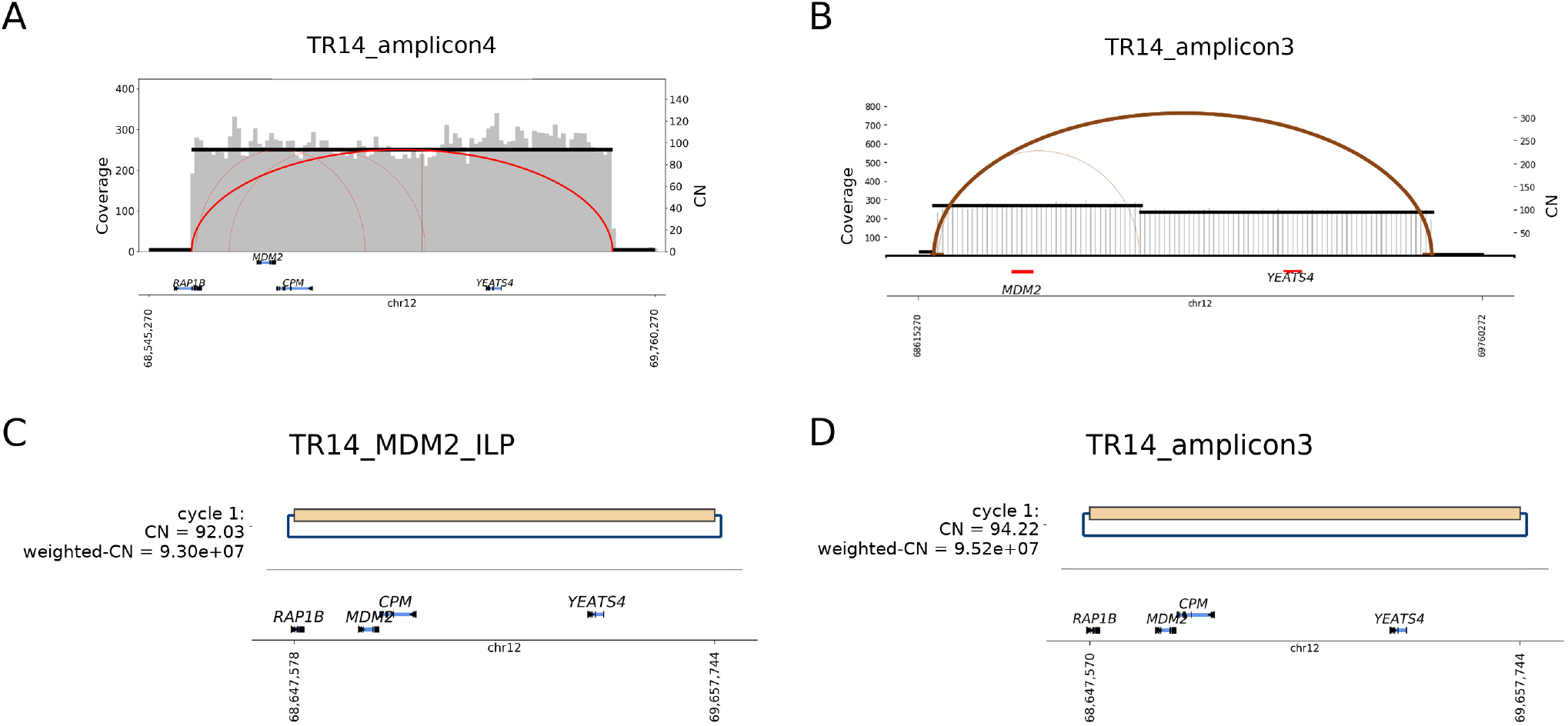
CE reconstruction on TR14 cell line, *MDM2* amplicon. (A) Long read amplicon graph. (B) Short read amplicon graph. (C) The heaviest cycle from CE reconstruction using the long-read graph, which is identical to CoRAL. (D) The heaviest cycle from CE reconstruction using the long-read graph, which is identical to AA.

**Supplemental Figure S9:**
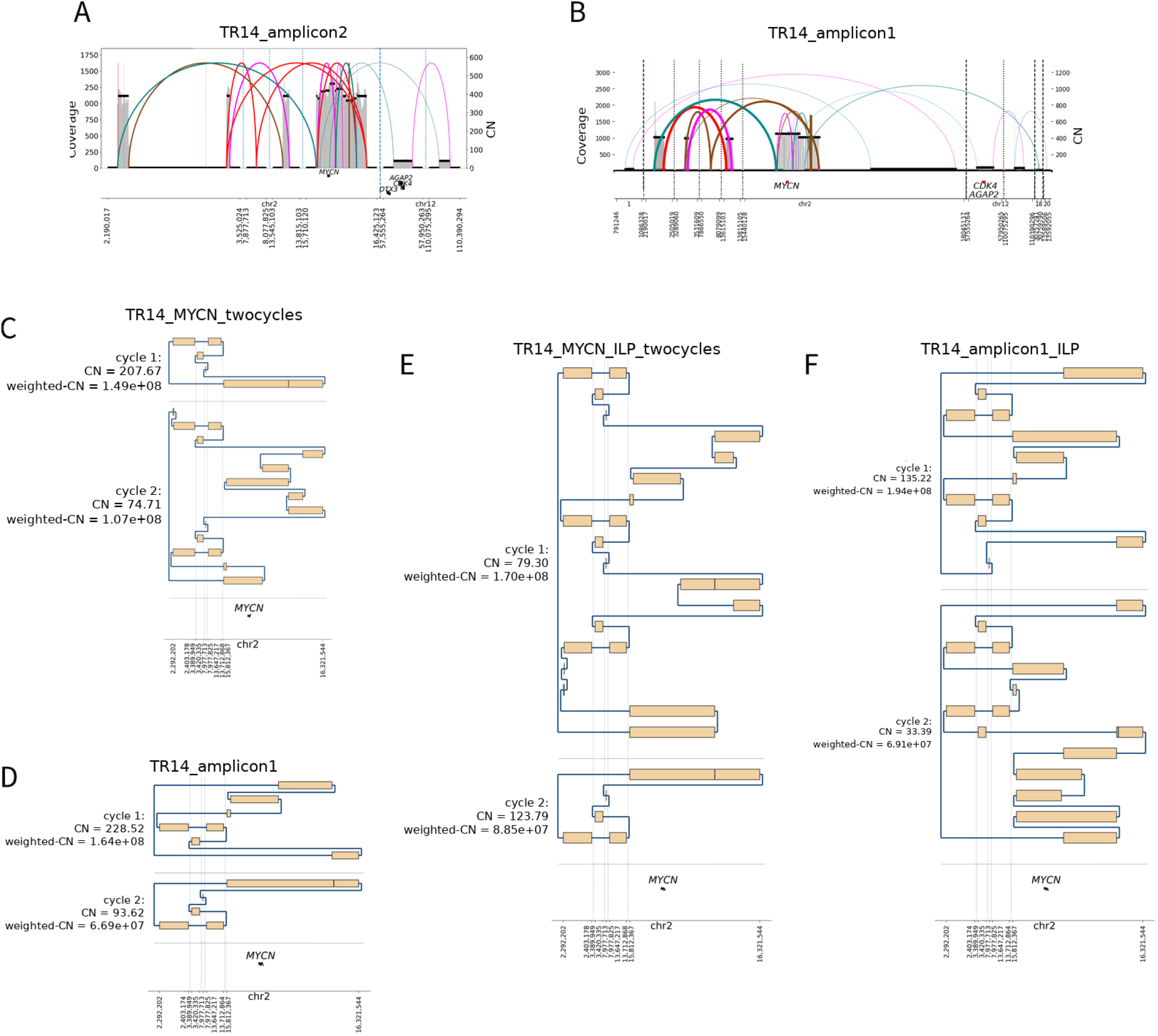
CE reconstruction on TR14 cell line, *MYCN* amplicon. (A) Long read amplicon graph. (B) Short read amplicon graph. (C) 2 heaviest cycles from CoRAL reconstruction. (D) 2 heaviest cycles from AA reconstruction. (E) 2 heaviest cycles from CE reconstruction, using long read graph. (F) 2 heaviest cycles from CE reconstruction, using short read graph.

**Supplemental Figure S10:**
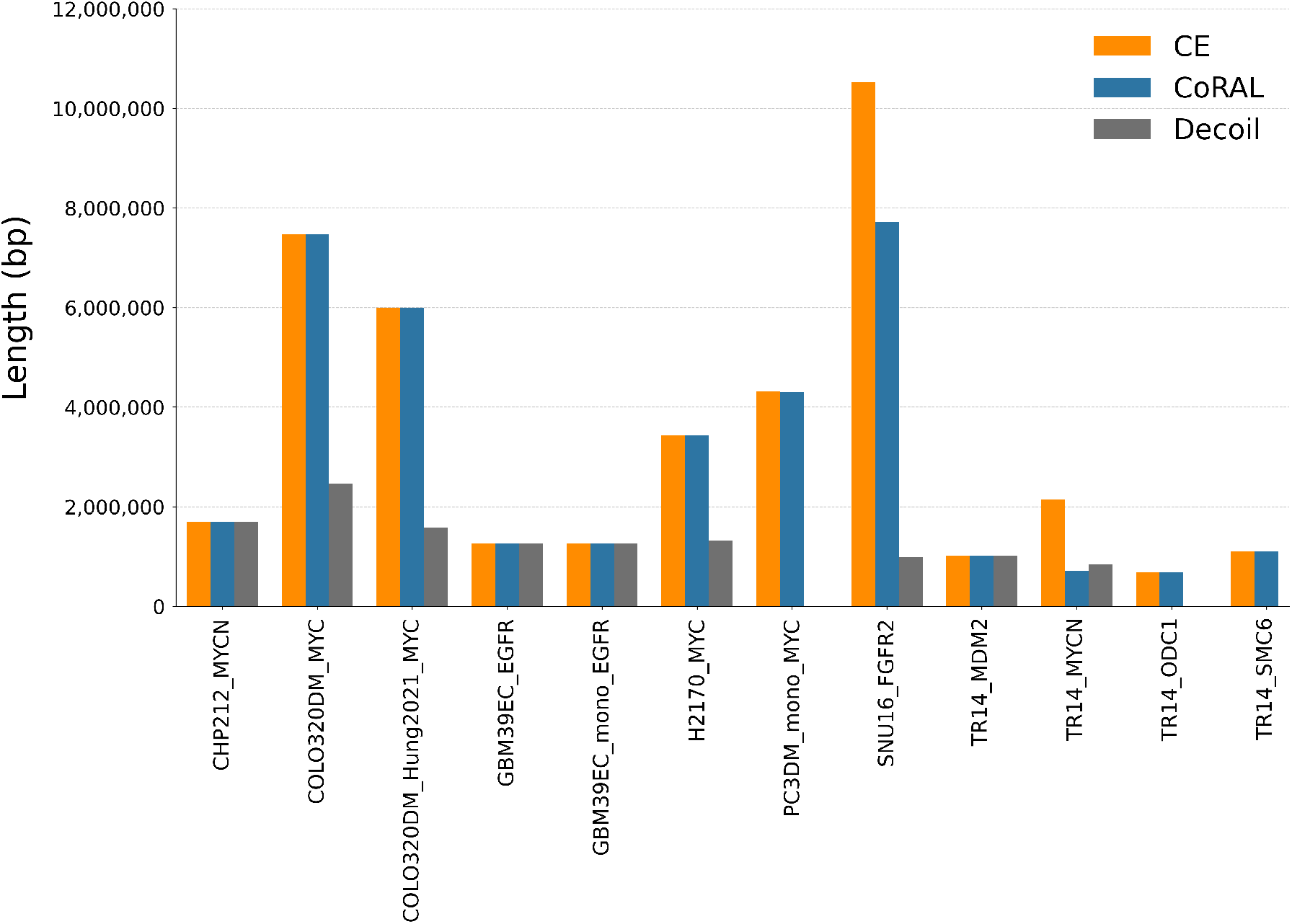
Length of the heaviest reconstructed cycle in long-read cancer cell lines. For each amplicon, the length of the heaviest cycle reconstructed by CE, CoRAL and Decoil. Decoil did not reconstruct any cycle for PC3DM mono *MYC*, TR14 *ODC1*, or TR14 *SMC6*. In several amplicons, including COLO320DM *MYC*, COLO320DM Hung2021 *MYC*, H2170 *MYC*, and SNU16 *FGFR2*, the Decoil cycle is shorter than the corresponding CE and CoRAL cycles.

**Supplemental Figure S11:**
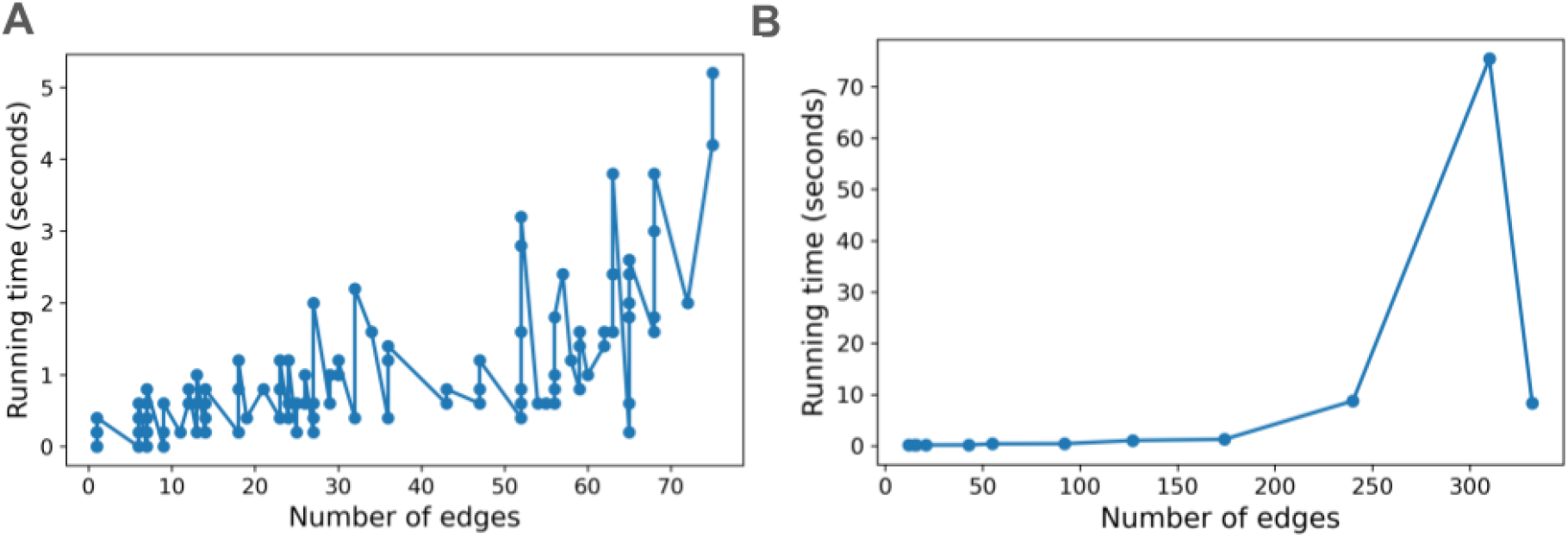
Running time vs the number of edges of the graph. (A) CE running time vs number of edges for simulated long-read data from cell lines. (B) CE running time for long-read (ONT) data on cell lines.

**Supplemental Figure S12:**
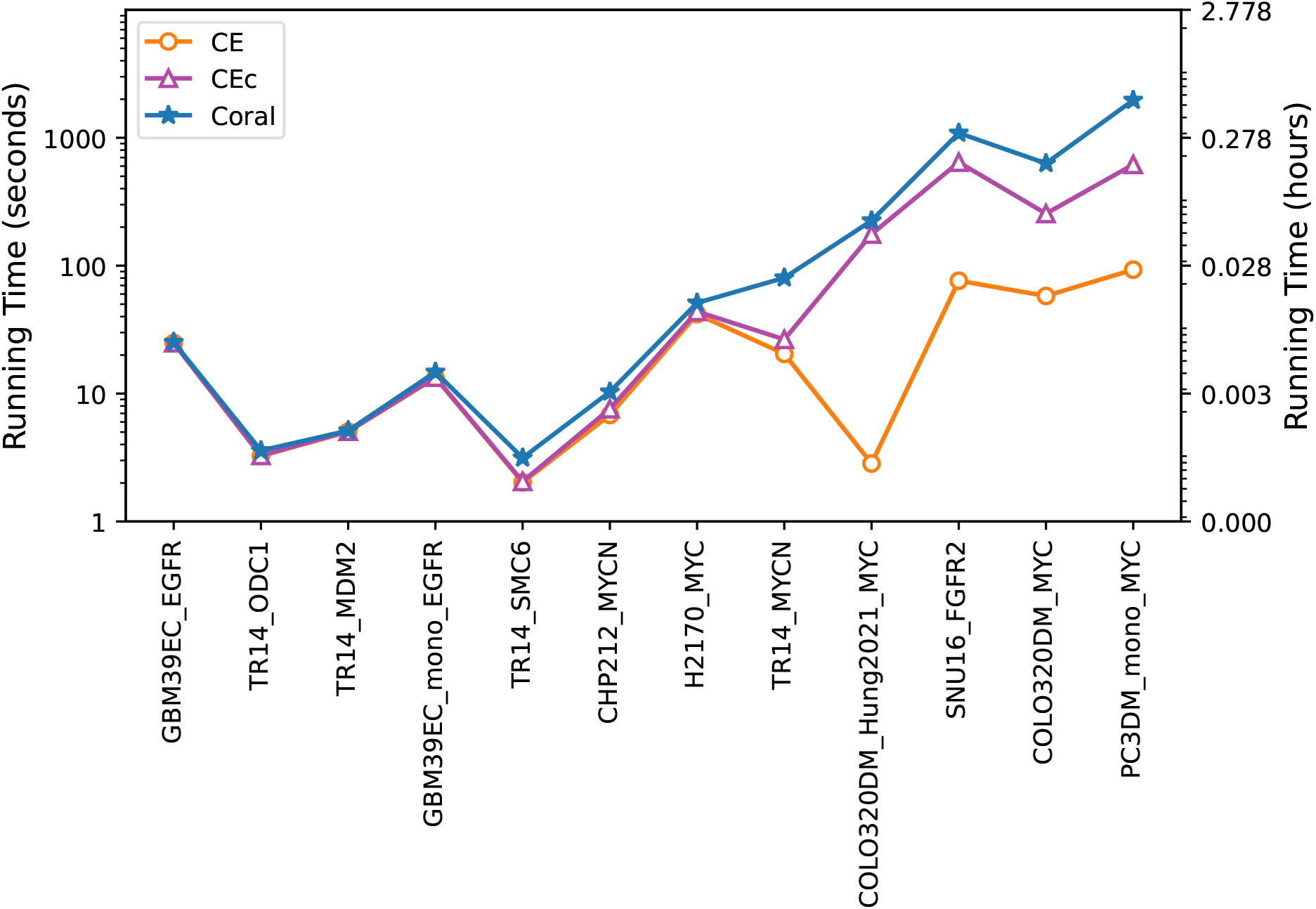
Running times of different methods for graph construction and cycle extraction. On ONT samples. Since CE and CEc take the breakpoint graph constructed by CoRAL as input, the same graph construction time is included for all three methods.

**Supplemental Figure S13:**
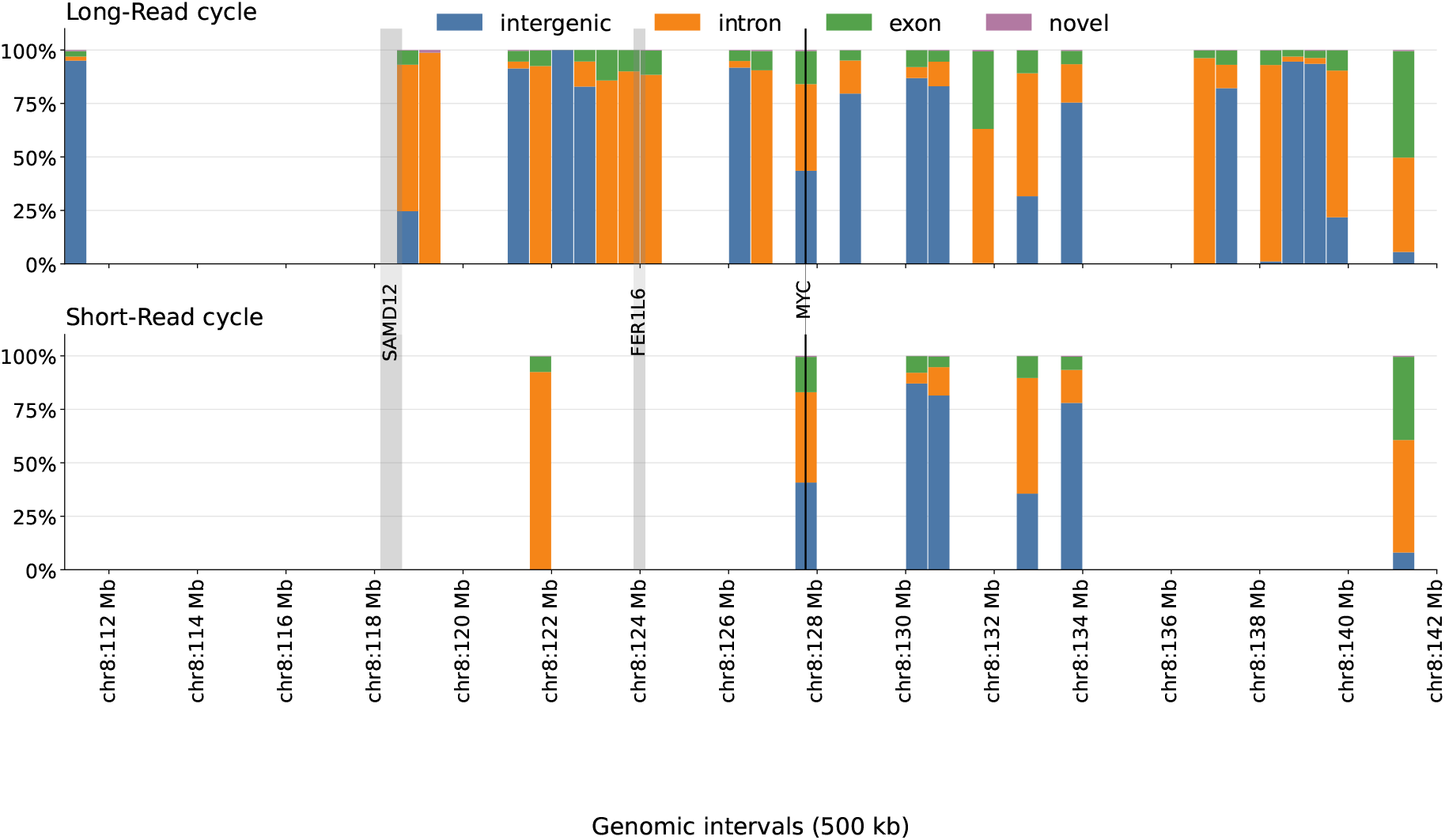
Comparison of gene-context annotation between the long-read and short-read PC3 ecDNA reconstructions. Each reconstructed cycle is partitioned into 500kb genomic intervals and annotated according to gene context using KaryoScope. Stacked bars show the fraction of each interval classified as intergenic, intron, exon, or novel sequence. Vertical gray bands indicate the locations of *SAMD12, FER1L6*, and *MYC*. The long-read reconstruction spans substantially more genomic sequence than the short-read reconstruction while preserving the overall gene-context composition and recovering additional genic regions.

**Supplemental Figure S14:**
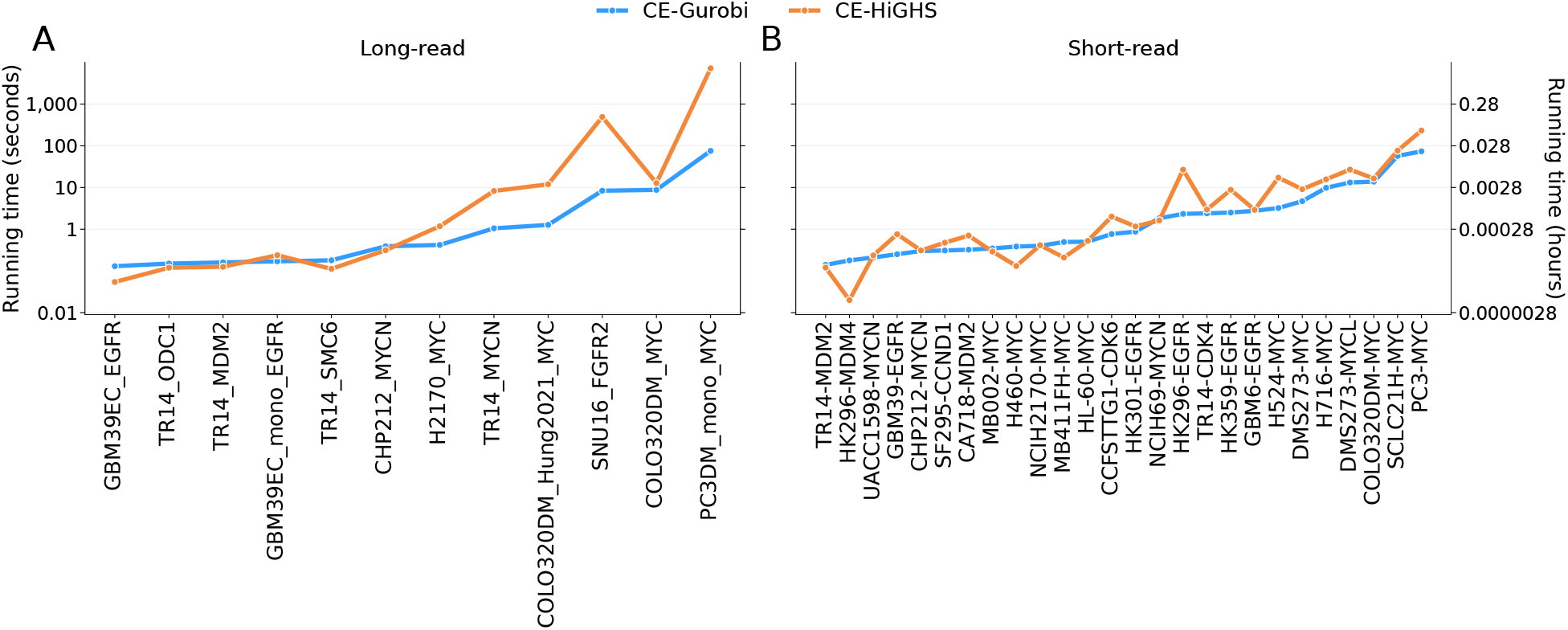
Running times of CE using Gurobi and HiGHS. (A) On ONT samples. (B) On Illumina samples. CE solves the same MILP using both solvers, so the difference only reflects the choice of solver.

## Supplementary Methods

### Full ILP for the *optimization* step

The full CE MILP formulation is provided below.

#### Parameters

*M* : maximum multiplicity allowed for concordant and discordant edges

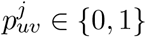: indicates if edge (*u, v*) is in subwalk *p*_*j*_

#### Variables

*F* ∈ ℝ : copy number assigned to edges in the solution but with multiplicity one.

*x*_*uv*_ ∈ {0, 1} : *x*_*uv*_ = 1 if edge (*u, v*) is selected, and 0 otherwise.

*f*_*uv*_ ∈ ℝ : copy number assigned to edge (*u, v*) in the solution, should be multiples of *F* .

*P*_*j*_ ∈ {0, 1} : indicates if subwalk *p*_*j*_ is satisfied 1 ≤ *j* ≤ *p*.

*d*_*u*_ ∈ {0, 1, 2, ..} : indicates the order of the nodes, used in connectivity constraints.

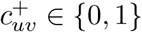: be a binary variable that is 1 if the direction is from *u* to *v*, again used in connectivity constraints.

##### CE MILP

CE maximizes the sum of LWCN and *γ*-weighted subwalk constraint using

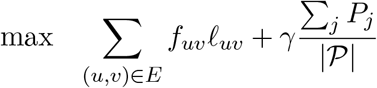

while satisfying the following constraints.

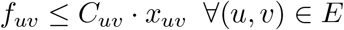
2. **balance**: The sum of the solution copy number of discordant and concordant edges that connect to a node is equal to the solution copy number of the sequence edge connected to that node.

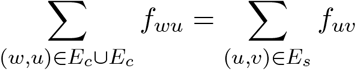
3. **copy number** *F* **with multiplicity one**: Note that if edge (*u, v*) is picked in the solution (i.e., *x*_*uv*_ = 1), then *F* ≤ *f*_*uv*_, but otherwise (*u, v*) imposes no constraint. For this purpose we introduce a sufficiently large constant *C*_max_ (e.g., *C*_max_ = *M* · max_(*u,v*)∈*E*_ *f*_*uv*_).

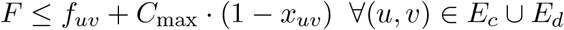
4. **discordant edge multiplicity constraint**: The multiplicity of a discordant edge is managed using

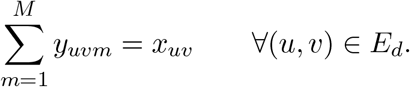

Subsequently the copy of number of the edge is given by the (quadratic) constraint

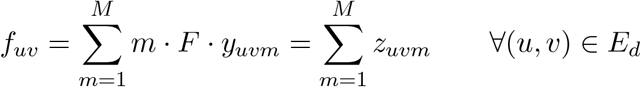

which guarantees *f*_*uv*_ are multiples of *F* . We linearize the quadratic constraints using

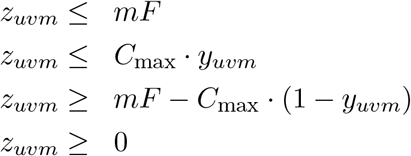
5. **subwalk constraints**: If a subwalk is in the solution (*P*_*j*_ = 1), all of its edges are in the solution. If a subwalk in not in the solution (*P*_*j*_≠ 0), there is not constraint here.

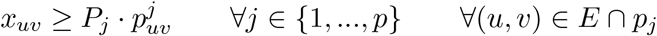
6. **connectivity constraints 1 (start node):** Assign *s* to be the start node. The order of *s* is 1.

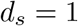

No direction into *s*.

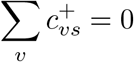

Direction out of s

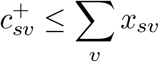
7. **connectivity constraints 2 (order of nodes):** If node *u* is not in the solution, then *d*_*u*_ = 0, otherwise *d*_*v*_ is a positive integer which reflects the order of the node.

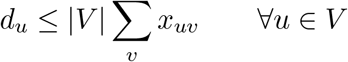
8. **connectivity constraints 3 (edge direction):** Only one direction from each edge in the solution

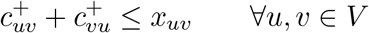
9. **connectivity constraints 4 (incoming edge for nodes):** If a node is part of the solution, and it is not *s* (the starting node), then it must have at least one incoming direction

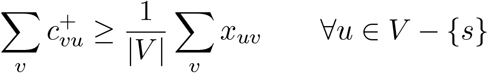
10. **connectivity constraints 5:**

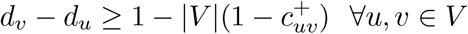

### Comparison of CE and CoRAL formulations

For completeness, we summarize and compare the formulations of CoRAL and CE. In CoRAL (specifically the max weight formulation), the objective function was quadratic. CoRAL comprised 20 classes of constraints; of which 4 were quadratic and the others were linear. We reformulated the same optimization problem with 16 classes of MILP constraints.

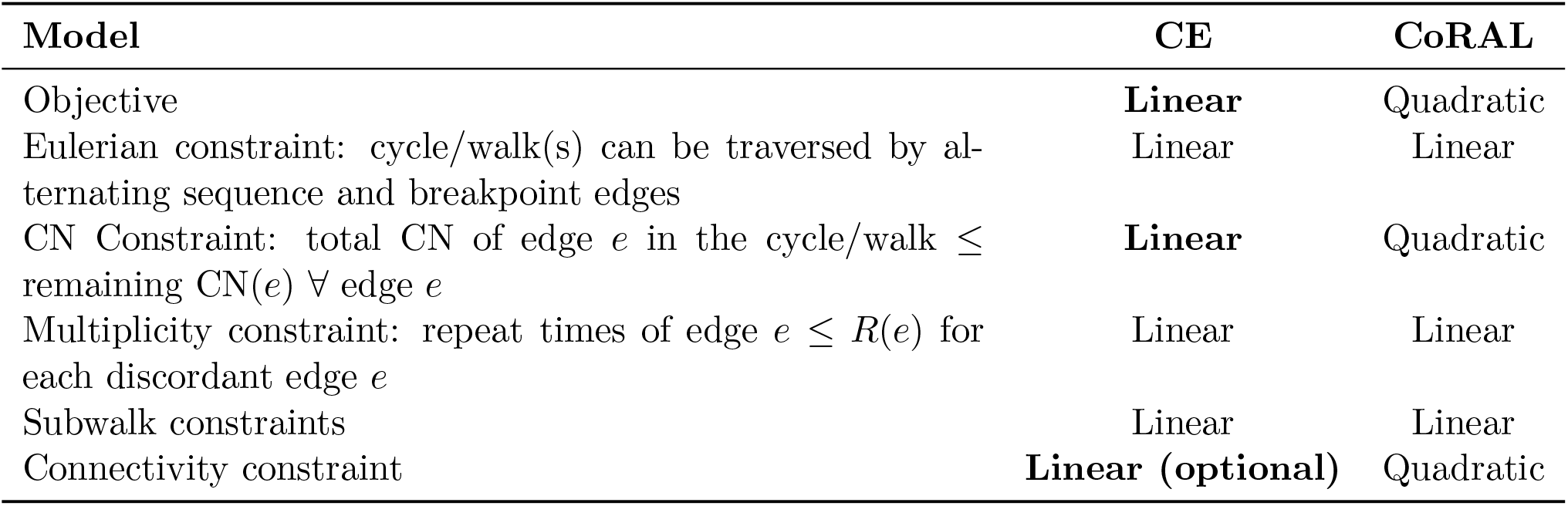

One key difference between CE and CoRAL is the treatment of edge multiplicities. In CE, we do not introduce an explicit multiplicity variable for edges. Instead, the model includes a variable representing the copy number of the ecDNA cycle, along with variables representing the copy number contributed by each edge. The multiplicity of an edge can then be calculated after solving the MILP optimization by dividing the edge copy number by the copy number of the ecDNA cycle. In contrast, CoRAL directly models edge multiplicities as integer variables within its optimization framework.

### Integer multiples of *F* of all edges in the solution

#### Lemma S1

*If the copy numbers of the discordant edges in the cycle are positive integer multiples of F (as enforced by Constraint 4), then the copy numbers of the concordant and sequence edges in the solution cycle are also positive integer multiples of F* .

*Proof*. We consider a canonical ordering of all nodes in the MILP solution based on their chromosomes and then on their nucleotide coordinates. By definition, node 1 in the solution is connected to a sequence edge and one or more discordant edges, but not to any concordant edge. We prove by induction that for each node *k* in the ordering,any edge incident on node *k* must have a copy number that is an integer multiple of *F* .

#### Base case (*k* = 1)

We know that node 1 is only incident on a single sequence edge, and all other edges connecting to 1 are discordant, with copy numbers that are multiples of *F* . By the balance condition, the sequence edge (1, 2) must also have a copy number that is an integer multiple of *F* . Therefore, all edges connected to node 1 have copy numbers that are positive integer multiples of *F* .

#### Induction hypothesis

Assume that every edge incident to nodes 1, 2, …, *k* has copy number equal to a positive integer multiple of *F* .

#### Inductive Step

We now consider node *k* + 1. All discordant edges incident on node *k* + 1 must have copy numbers that are multiples of *F*, and we need to investigate the concordant and sequence edges (denoted *e*_*c*_, *e*_*s*_, respectively) incident on *k* + 1. We have two cases.

First, suppose CN[*e*_*c*_] = 0. Then, by the balance condition, CN[*e*_*s*_] must be an integer multiple of *F* . Second, we consider the case CN[*e*_*c*_] *>* 0, then both nodes must be a part of the solution, and by our ordering criteria, *e*_*c*_ = (*k, k* + 1), connects node *k* to *k* + 1. Then, using induction hypothesis, CN[*e*_*c*_] must be an integer multiple of *F*, and by balance, CN[*e*_*s*_] must be a positive integer multiple of *F* .

### Implementation details

#### Preprocessing amplicon graphs

While CE accepts an amplicon graph directly from the output of AA (Deshpande et al. 2019) or CoRAL (Zhu et al. 2024), it replaces each foldback edge (*u, u*) ∈ *E*_*d*_ with (*u, u*_1_) ∈ *E*_*d*_, (*u*_1_, *u*_2_) ∈ *E*_*s*_, and (*u*_2_, *u*) ∈ *E*_*d*_, and setting 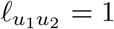. This construction allows alternate paths in the model to traverse through foldbacks and, by assigning a positive length, encourages their inclusion in the solution based on the objective function.

#### Connectivity enforcement cycle extraction

When CEc aims to extract a cycle (to avoid multiple disconnected cycles), it connects artificial source (*s*) and sink (*t*) nodes to all nodes in the graph. To extract exactly one cycle, CEc finds a path from *s* to *t*, with *s* and *t* connected to the same node in the solution. To ensure feasibility with respect to the alternating edge constraints, *s* is connected to other nodes via discordant edges, while *t* is connected via sequence edges. The copy number assigned to each of these newly added edges equals the maximum copy number of the edges originally adjacent to that node. The length of the sequence edge connecting *t* is set to 1 - positive, but short - so that it encourages selection in the solution.

#### Extract paths from graph

The CE(c) method is an iterative framework that, in each iteration, takes as input an amplicon graph with copy number annotated edges. It formulates and solves a mixed-integer linear programming (MILP) problem to identify a subset of edges, each with an assigned copy number, that can form a cycle. After each iteration, the graph is updated by subtracting the copy numbers of the selected edges from their corresponding values in the graph. The updated graph is then used as input for the next iteration to extract additional cycles. Although CE/CEc can also extract paths, it prioritizes the detection of cycles. Once no cycles remain, the algorithm proceeds to extract paths.

If CE/CEc aims to extract a path after all cycles have been extracted, it introduces artificial source (*s*) and sink (*t*) nodes. Both nodes are connected to all other nodes in the graph via discordant edges, and each new edge is assigned a copy number equal to the maximum copy number of the edges originally adjacent to that node.

### Cell lines used in this paper

For building the amplicon graph, we used the following: AmpliconArchitect (AA) version 1.5.r1; CoRAL version 2.2.0.

### Metaphase FISH

Half a million PC3-DM5 cells were seeded on a 6mm dish followed by a 4-hour KaryoMAX Colcemid (Gibco) treatment at 100ng mL^−1^ to arrest cells into metaphase. Cells were then trypsinized, washed in 1X PBS, followed by a 20-min 0.075M KCl hypotonic buffer incubation at 37°C. Freshly prepared Carnoy’s fixative (3:1 methanol:glacial acetic acid) was added to the cells, fully resuspended by pipetting and spun down. Two additional washes in Carnoy’s fixative were performed, and the cell pellet was resuspended in Carnoy’s fixative. Cells were dropped onto a humidified glass slide and allowed to air-dry. The slide was briefly equilibrated in 2X SSC buffer, followed by dehydration in ascending ethanol concentrations of 70%, 85% and 100% for 2 mins each. DNA FISH probes (Empire Genomics) were diluted at 1:10 in hybridization buffer. The following FISH probes were used to detect *MYC* (RP11-440N18), *SAMD12* (RP11-671M23) and *FER1L6* (RP11-1112G17). The probe mixture was added to the sample and covered with a 22×22 coverslip. The sample was subjected to a 3-min heat denaturation at 75°C, followed by an overnight hybridization at 37°C in a humidified slide moat. The next day, the coverslip was carefully removed from the glass slide and the sample was washed in 0.4X SSC for 2 mins, followed by another 2-min wash with 2X SSC containing 0.1% Tween 20. The slide was incubated with 2X SSC containing DAPI (50 ng mL^−1^) for another 10 mins to stain DNA content. Followed by a brief ddH2O wash, the sample was mounted with ProLong Diamond Antifade and air-dried. Images were acquired on a Leica DMi8 widefield microscope with a 63x oil objective.

## Footnotes

1 While CE prioritizes cyclic walks to reconstruct ecDNA, it can also identify non-cyclic walks when no heavy cycles can be found (see Supplementary Methods for details).

